# Distribution and strength of interlaminar synaptic connectivity in mouse primary visual cortex revealed by two-photon optogenetic stimulation

**DOI:** 10.1101/2019.12.13.876128

**Authors:** Travis A. Hage, Alice Bosma-Moody, Christopher A. Baker, Megan B. Kratz, Luke Campagnola, Tim Jarsky, Hongkui Zeng, Gabe J. Murphy

## Abstract

The most common synaptic connections between neurons in different cortical layers form the basis of the canonical cortical microcircuit. Understanding of cortical function will require further development and application of methods to efficiently characterize synaptic connections within and outside of the canonical pathway. Accordingly, we measured synaptic inputs onto superficial excitatory neurons in response to sequential two-photon optogenetic stimulation of neurons in deeper layers. Layer 4 excitatory neurons and somatostatin-neurons within layer 2/3 represented the most common sources of input. Although connections from excitatory and somatostatin-neurons in layer 5 were less common, the amplitudes of synaptic responses were equally strong. We examined synaptic strength across all connections, as well as the relationships between the strength of connections diverging from a common presynaptic neuron or converging to a single target. While the overall distribution indicates synaptic weight is concentrated to a few connections, strong excitatory connections are distributed across cells.

## Introduction

The receptive field properties of visual cortical neurons are governed by long-range synaptic connections formed across brain areas and local inputs formed between cells within a cortical area. Lack of detailed information about both long-range and local - i.e., within layer/horizontal connections and translaminar connections - limits mechanistic understanding of how sensory information is propagated and transformed; so, too, does the absence of detailed information about the patterns of convergence of synaptic inputs to neurons in the recipient layers and divergence of outputs from the projecting layers.

L2/3 pyramidal neurons represent a critical node in the canonical pathway by which thalamic input is conveyed to L4 neurons in a primary cortical area, on to L2/3 and then L5 neurons in the same area, and ultimately on to higher cortical areas. Although L2/3 neurons in V1 are known to receive direct input from dorsal lateral geniculate nucleus (dLGN)(Ji et al., 2016; Morgenstern et al., 2016; D’Souza et al., 2019) and from higher visual areas (DSouza et al., 2016; Young et al., 2019), monosynaptic rabies tracing suggests the majority of presynaptic inputs arise from neurons within V1 (Liu et al., 2013; Wertz et al., 2015). These local inputs include well-established recurrent excitatory connections between L2/3 pyramidal neurons, horizontal inhibition from nearby interneurons, and ascending inputs from L4 excitatory neurons that carry sensory information from thalamus.

Local ascending projections from neurons in layers 5 and 6 to L2/3 have been described in anatomical studies of sensory cortex and are thought to represent feedback from later stages of cortical processing (Burkhalter, 1989; Yoshioka et al., 1994). Experiments performed in juvenile visual and somatosensory cortex using laser scanning photostimulation combined with glutamate uncaging have identified connections from excitatory neurons in L5 to L2/3 (Dantzker and Callaway, 2000; Bureau et al., 2004; Shepherd et al., 2005; Shepherd and Svoboda, 2005; Yoshimura et al., 2005; Hooks et al., 2011; Xu et al., 2016). However, connections from L5 to L2/3 have been scarce or absent, when probed via simultaneous electrophysiological recordings in rodent barrel or visual cortex (Thomson and Bannister, 1998; Reyes and Sakmann, 1999; Thomson et al., 2002; Lefort et al., 2009; Jiang et al., 2015). The L5 to L2/3 connection is most often observed from L5a and therefore thought to originate from IT-type L5 pyramidal neurons (Dantzker and Callaway, 2000; Bureau et al., 2004; Shepherd et al., 2005; Shepherd and Svoboda, 2005; Yoshimura et al., 2005; Hooks et al., 2011; Xu et al., 2016). It was recently demonstrated that feedback projections from anteromedial or lateromedial visual areas preferentially target IT-type L5 neurons in V1 to generate loops between higher and lower visual areas (Young et al., 2019), raising the possibility that local ascending projections from L5 could transmit behavioral or brain-state-related information to superficial layers. A direct comparison of the translaminar excitation from layers 4, 5 and 6 to L2/3 in mature rodent V1 has not yet been made and will be necessary to understand the contributions of feedforward- and feedback- inputs to the response properties of L2/3 neurons.

Likewise, direct comparisons of horizontal and vertical inhibition are needed to understand the role of local connectivity in shaping neuronal responses. Intralaminar inhibition from interneurons to nearby pyramidal cells has been described as abundant and non-selective (Packer and Yuste, 2011; Fino et al., 2013; Karnani et al., 2014). Such horizontal inhibition from somatostatin (Sst) interneurons shapes the receptive field properties of nearby pyramidal neurons in L2/3 of mouse visual cortex (Adesnik et al., 2012). Several patterns of vertical inhibition (mediated by translaminar inhibitory connections) have been described for Sst-interneurons (Kapfer et al., 2007; Silberberg and Markram, 2007; Jiang et al., 2015; Anastasiades et al., 2016; Naka et al., 2019) as well as parvalbumin-interneurons (Olsen et al., 2012; Bortone et al., 2014).

Furthermore, subtypes of Sst-interneurons may be differentially integrated into distinct layer-specific subnetworks (Muñoz et al., 2017; Naka et al., 2019). Although intra- and inter-laminar inhibitory inputs to L2/3 of V1 have been compared via one-photon photostimulation activation of GABAergic interneurons via either caged-glutamate (Xu et al., 2016) or channelrhodopsin-2 expression (Kätzel et al., 2011), potentially unique roles of subsets of GABAergic interneurons may be revealed by photostimulation techniques with increased genetic and spatial specificity.

In this study, we measured the probability, spatial distribution and strength of translaminar synaptic connectivity by combining multicellular patch-clamp recording (up to 4 postsynaptic neurons) with two-photon optogenetic stimulation of putative presynaptic neurons. Combining multicellular recording with multiphoton photostimulation permitted high-throughput characterization of synaptic connectivity (517 total connections identified from 7968 probed) and allowed us to measure relatively sparsely connected elements of the cortical microcircuit. Consistent with the canonical microcircuit, L4 excitatory neurons and Sst neurons within L2/3 displayed the highest rates of connectivity onto L2/3 pyramidal neurons. Additionally, we observed relatively sparse connections from excitatory neurons in L5 and Sst neurons in layers 4 and 5. The strength of both excitatory and inhibitory synaptic connections onto L2/3 pyramidal neurons resembled a lognormal distribution, as described for connections between other classes of cells (Sayer et al., 1990; Feldmeyer et al., 2002; Song et al., 2005; Lefort et al., 2009). Although less common, the amplitudes of excitatory and inhibitory postsynaptic potentials (EPSPs and IPSPs) from cells in L5 were similar to the more canonical excitatory connections from L4 and intralaminar inhibitory connections. Use of multicellular recording, allowed us to identify individual presynaptic cells with divergent connections onto multiple L2/3 pyramidal cells. Diverging outputs from Sst neurons were most often confined to postsynaptic cells within 100 μm of each other and the amplitudes of Sst connections that shared a presynaptic source were moderately correlated. By contrast, connections from individual excitatory neurons in L4 or L5 diverged to target cells in L2/3 separated by hundreds of microns, and the strengths of diverging translaminar excitatory connections were not correlated. Whereas we see evidence that some Sst neurons more strongly inhibit their local network than others, strong translaminar excitatory connections from appear to be more evenly distributed across presynaptic and postsynaptic cells.

## Results

We measured translaminar synaptic connectivity and strength in primary visual cortex of mature mouse via two-photon stimulation of opsin-expressing neurons. We used layer-specific Cre lines - Scnn1a and Rorb-Cre, Tlx3-Cre, and Ntsr1-Cre to stimulate excitatory neurons in cortical layers 4,5, and 6, respectively. Sst-Cre was used to stimulate Sst-interneurons.

### Characterization of two-photon stimulation of ChrimsonR-expressing neurons

A number of methods have been developed for two-photon stimulation of neurons, with varying trade-offs in terms of reliability, spatial specificity, ability to stimulate multiple neurons, and complexity (Emiliani et al., 2015). We pursued rapid galvo-based spiral scanning of opsin-expressing neurons (Rickgauer and Tank, 2009; Prakash et al., 2012; Packer et al., 2012) as a method to stimulate ChrimsonR-expressing neurons (Klapoetke et al., 2014). We tested the reliability and specificity of two-photon stimulation using two methods of ChrimsonR expression. The first utilized the Ai167 TIGRE2.0 line, which we previously found generated robust one-photon-evoked photocurrents (Daigle et al., 2018). The second method took advantage of recent evidence that the addition of short peptide sequences restricts opsin trafficking to the soma and proximal of dendrites of neurons, thereby improving the spatial specificity of photostimulation (Baker et al., 2016; Shemesh et al., 2017). Therefore, we utilized a combinatorial approach in which we performed stereotaxic injections of a Cre-dependent adeno-associated virus (AAV) that expresses soma-targeted-ChrimsonR (flex-ChrimsonR-EYFP-KV2.1) into primary visual cortex of Cre-positive mice.

We used loose-seal cell attached recordings to minimally disturb intrinsic cell properties when measuring light-evoked generation of action potentials (APs) (Figure 1A). Across all cells, the minimum power necessary to generate one or more APs in ChrimsonR-expressing neurons ranged from 3.4-94 mW (10 ms photostimulus duration, measured from 7-19 neurons for each Cre line/opsin combination, Figure 1B). Although the distributions vary somewhat between Cre lines, we observed spiking in 90-100% of targeted cells in all Cre lines when using an 85 mW photostimulus. AAV-transfected neurons in the Sst line were very sensitive to photostimulation, with half of cells generating APs in response to 8 mW photostimuli (Figure 1B). Based on these results, we used 85 mW as our primary photostimulation power for probing synaptic connectivity in experiments using Ai167 and excitatory Cre lines transfected with AAV. To avoid off-target activation of the more photosensitive AAV-transfected Sst-neurons, we used a lower stimulus intensity of 35 mW.

**Figure 1.**
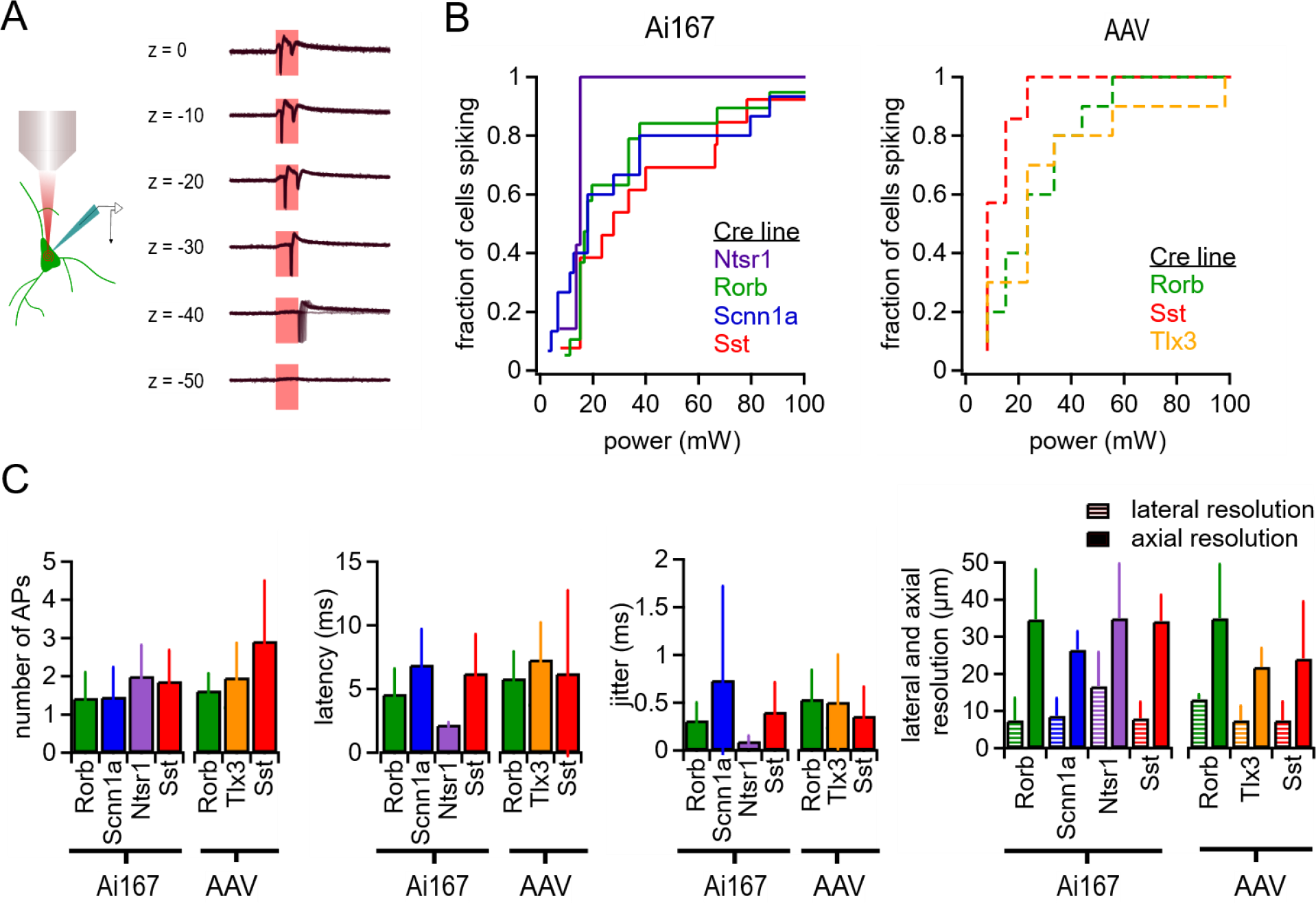
Characterization of two-photon evoked spiking. **(A)** Examples of APs evoked by two-photon stimulation of a Rorb:Ai167 neuron with indicated axial offsets. Each panel contains 10 overlaid sweeps. **(B)** Cumulative probability of light-evoked spiking versus photostimulus power for the indicated Cre lines crossed to Ai167 or injected with AAV expressing soma-targeted ChrimsonR. **(C)** The average number of APs evoked per photostimulus, latency, jitter, and spatial resolution of photostimulation (measured using the same power used for mapping experiments) across cells for all Cre-line and opsin expression strategies used in this study. Error bars represent standard deviation. Minimum power, latency and jitter data were collected from a total of 88 neurons (7-19 neurons per Cre line-opsin combination). Lateral resolution was measured from a total of 51 neurons (4-12 per Cre line-opsin combination. Axial resolution was measured from a total of 56 neurons (5-13 per Cre line-opsin combination).

We measured the number of light-evoked APs, the latency to the first AP and the associated temporal jitter (standard deviation of the latency) as a function of stimulus intensity (Figure 1 - Supplement 1). The mean and standard deviation of these values at the powers used for mapping experiments are presented in Figure 1C. Latency and jitter decreased with higher stimulus intensities, although the effect on jitter was less dramatic as many cells displayed sub-millisecond jitter even with low power stimuli. Some cells generated multiple APs when stimulated (Figure 1 B,C and Figure 1-Supplement 1). Similarly, we observed instances of multiple light-evoked postsynaptic potentials (PSPs) in subsequent mapping experiments (for example see Figure 3A) and utilized exponential deconvolution to aid in identifying such cases (see Methods, Figure 5 - Supplement 1).

Finally, we characterized the spatial resolution of photostimulation. Consistent with previous reports (Rickgauer and Tank, 2009; Prakash et al., 2012; Packer et al., 2012), and the more axially elongated point spread function of two-photon excitation, the probability of light-evoked spiking decreased rapidly with lateral offset of the photostimulus target (P_spiking_ < 0.5 at 7-17 μm; Figure 1C, Figure 1 Supplement) whereas greater axial offset was necessary to observe the same decrease in light-evoked spiking (P_spiking_ < 0.5 at 22-35 μm; Figure 1C, Figure 1-Supplement 1). The results demonstrate rapid scanning of ChrimsonR-expressing neurons can reliably activate multiple classes of neurons, but also reveal some differences in sensitivity across Cre lines and methods of expression. We accounted for this in our photostimulation paradigms and wish to highlight the need for careful measurement of presynaptic light-evoked spiking even when using a common means of expression and photostimulation.

### Translaminar connectivity from excitatory neurons measured using two-photon stimulation

The excitatory synaptic connection from L4 neurons to L2/3 pyramidal cells represents the primary feedforward projection within the cortex (immediately following input from the thalamus to L4 cells). To probe physiological features of this connection we recorded from L2/3 pyramidal cells in mice in which ChrimsonR was expressed principally in excitatory neurons of L4 (Rorb-Cre and Scnn1a-Cre mice). We measured synaptic connections from Rorb neurons to L2/3 pyramidal cells using both transgenic ChrimsonR expression and AAV-mediated expression of soma-targeted ChrimsonR. Although both Cre lines sparsely label neurons outside of L4 - primarily in L5 - the spatial specificity provided by two-photon stimulation allows sequential and independent stimulation of individual cells.

An example experiment using a Rorb:Ai167 mouse is shown Figure 2A. In this experiment we measured input to a single L2/3 pyramidal neuron from 43 putative presynaptic Rorb-labeled neurons; traces represent a connected Rorb cell in L4 (photostimulus 1), a neighboring non-connected L4 neuron (photostimulus 2), and a connected Rorb neuron in L5 (photostimulus 3). Nearly all identified presynaptic partners to L2/3 pyramidal neurons in L4 were within 300 μm of lateral distance from the postsynaptic cell (Figure 2B). Within this distance connection rates measured using Rorb:Ai167, Rorb:AAV or Scnn1a:Ai167 were between 7.1% - 10.6% and were not significantly different (Chi-squared p=0.09; Rorb:Ai167: 10.6%, 93/880 probed, Rorb:AAV: 7.1% 32/450 probed, Scnn1a:Ai167: 7.4%, 11/148 probed).

**Figure 2.**
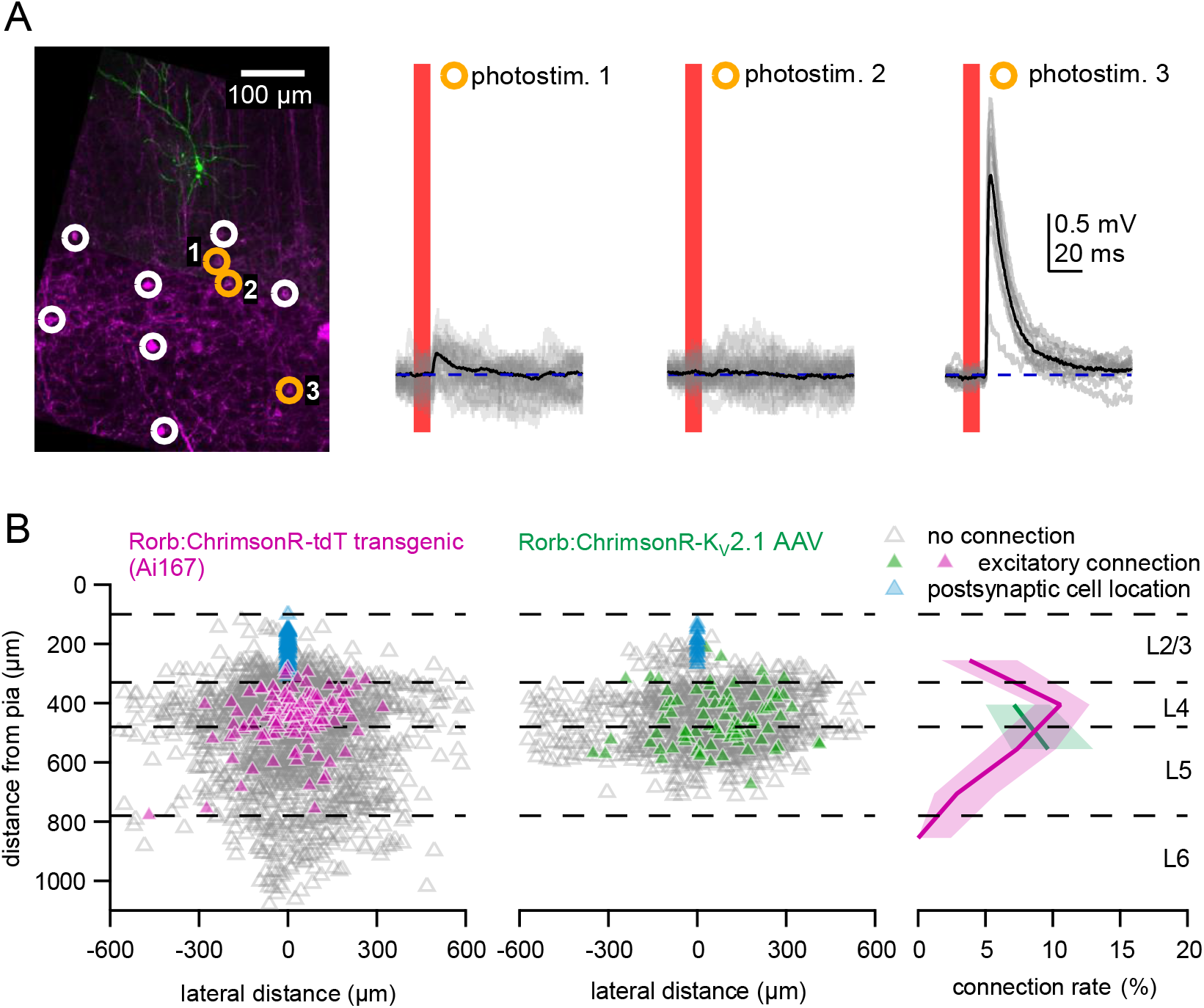
Measurement of excitatory connectivity from Rorb+ neurons in L4 and L5 to L2/3 pyramidal cells. **(A)** Example experiment measuring connectivity from L4 and L5 excitatory neurons labeled in acute slice from a Rorb:Ai167 mouse. *Left*: Flattened z-stack of recorded L2/3 pyramidal neuron filled with Alexa 488 (green) and td-Tomato signal from ChrimsonR-expressing neurons (magenta). Yellow circles indicate photostimulated cells with responses plotted in right panels. Responses following stimulation of cells in white circles are not shown. *Right*: Responses to photostimulation. Photostimulus occurred during the red bar, black lines are an average of individual sweeps (grey). **(B)** Summary maps for all experiments with Rorb-Cre mice crossed to Ai167 (*left*) or injected with AAV encoding soma-targeted ChrimsonR (*center*). Locations of neurons are plotted according to their distance from the pia and the lateral distance between putative pre- and postsynaptic cells with the following color scheme: *blue:* postsynaptic (recorded) L2/3 pyramidal cells *grey*: photostimulated neurons that did not evoke a synaptic response in the recorded cell. *magenta/green*: Stimulated cells that produced an EPSP using Ai167 or AAV-mediated opsin expression. Dashed lines represent approximate layer boundaries. Positive lateral distances correspond to presynaptic cells posterior to the postsynaptic cell. *Right:* The connection rate of Rorb-labeled neurons to L2/3 pyramidal cells plotted against the presynaptic cell’s distance from the pia. Measurement of connection rate versus pia distance was limited to cells <300 μm lateral distance. Shading indicates 95% confidence intervals.

Excitatory projections from L5 to L2/3 have been described as tightly focused and weak (Thomson and Bannister, 1998; Reyes and Sakmann, 1999; Thomson and Bannister, 2003; Lefort et al., 2009; Jiang et al., 2015) relative to inputs from L4 (Douglas and Martin, 2004). In this context, we were surprised to find many connections to L2/3 pyramids from Rorb neurons located in L5 using either transgenic or AAV-mediated opsin expression (Figure 2B). Furthermore, the observed rate of connectivity from Rorb neurons located in L5 was only slightly lower than the rate of Rorb+ neurons in L4 (Figure 2B; combing Ai167 and AAV data within 300 μm lateral distance, Rorb L4: 125/1330, 9.4%, Rorb L5: 71/994, 7.1%, Fisher’s Exact p=0.06).

Many of the Rorb L5 connections were found in the superficial half of L5 (Figure 2B), leading us to ask if these connections arose from “misplaced” L4 Projection neurons. Alternatively, given that subsets of Rorb neurons resemble IT type L5 neurons in terms of transcriptomic profiles (Tasic et al., 2016) and long-range projection patterns (Harris et al., 2019), it is possible that ectopic Rorb-Cre expression in L5 IT neurons allowed us to identify local projections from L5 IT neurons (that are primarily associated with inter-areal projections). To more deliberately measure the local projections of L5 IT neurons, we used the Tlx3-Cre line (Kim et al., 2015) to drive soma-targeted ChrimsonR expression. An experiment illustrated in Figure 3A shows an example in which combining multicellular recording with photostimulation revealed divergence of connectivity from a L5 IT neuron (photostimulus 2) to multiple L2/3 pyramidal cells. Across experiments, we observed Tlx3 to L2/3 connectivity at a slightly lower, but statistically similar rate compared to L5 Rorb connections (Figure 3B, Tlx3 all data: 60 found of 1420 probed, 4.2%, Tlx3 within 300 μm lateral distance: 55 found of 1010 probed, Fisher’s Exact comparing Rorb L5 to Tlx3, within 300 μm lateral distance p=0.12). In total, L5 to L2/3 connectivity was assayed via transgenic ChrimsonR expression driven by Rorb-Cre, as well as soma-targeted ChrimsonR expression driven by either Rorb-Cre or Tlx3-Cre. All 3 experiments suggest L5 IT type neurons are a significant source of excitatory input to L2/3 pyramidal neurons in mature mouse V1.

**Figure 3.**
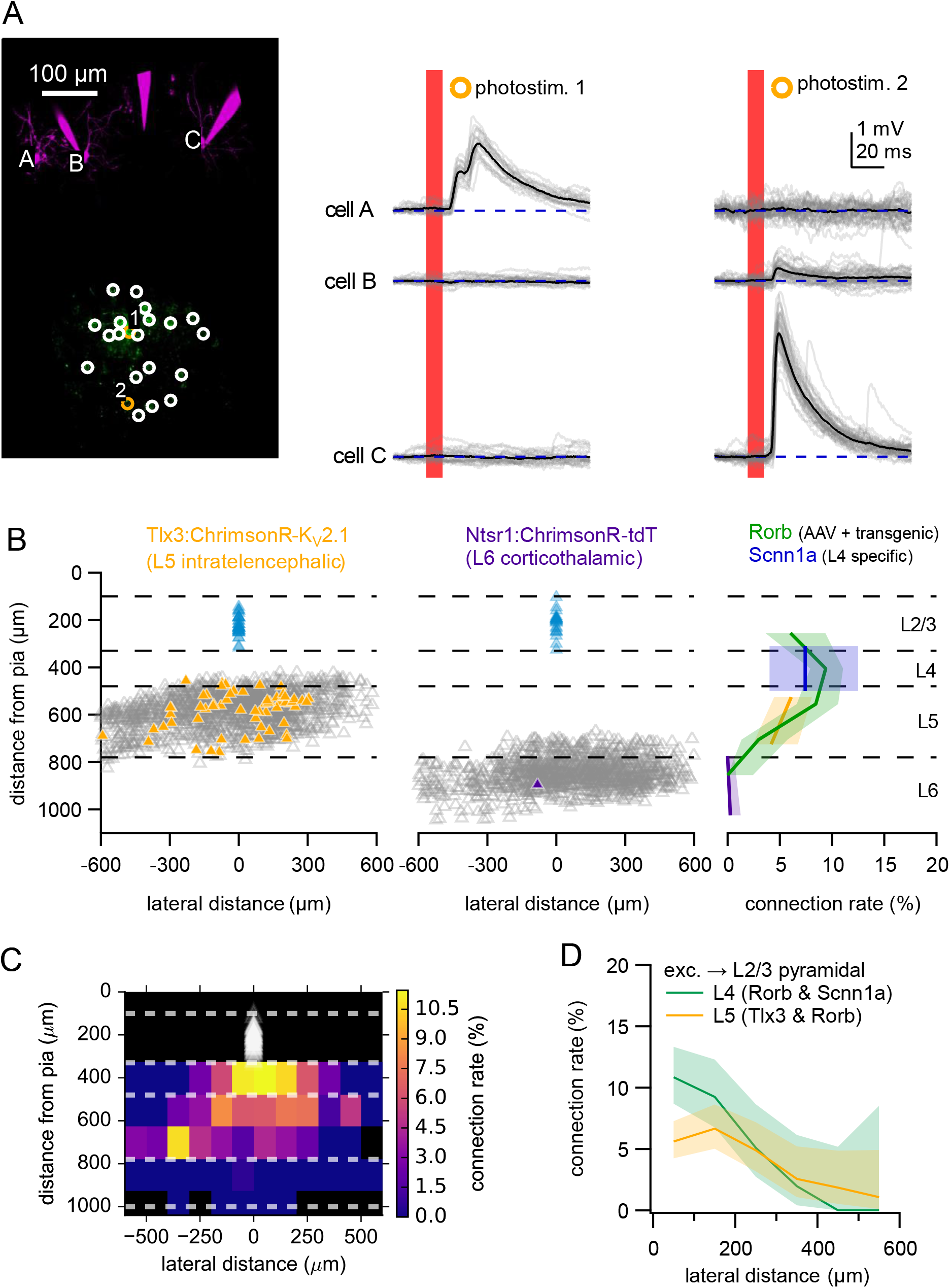
Measurement of excitatory connectivity from L5 and L6 to L2/3 pyramidal neurons. **(A)** Example experiment measuring connectivity from Tlx3+ excitatory neurons to L2/3 pyramidal neurons. *Left*: Flattened z-stack of recorded L2/3 pyramidal neurons filled with Alexa 594 (magenta) and EYFP signal from soma-targeted ChrimsonR-expressing neurons (green). Yellow circles indicate photostimulated cells with responses plotted in right panels. Responses following stimulation of cells in white circles are not shown. *Right*: Responses to photostimulation. Photostimulus occurred during the red bar, black lines are an average of individual sweeps (grey). **(B)** Summary maps for all experiments with Tlx3-Cre (*left*) or Ntsr1-Cre (*center*) mice. Locations of neurons are plotted according to their distance from the pia and the lateral distance between putative pre- and postsynaptic cells with the following color scheme: *blue:* postsynaptic (recorded) L2/3 pyramidal cells *grey*: photostimulated neurons that did not evoke a synaptic response in the recorded cell. *orange/purple*: Stimulated cells that produced an EPSP using Tlx3-Cre or Ntsr1-Cre. Dashed lines represent approximate layer boundaries. Positive lateral distances correspond to presynaptic cells posterior to the postsynaptic cell. *Right:* The connection rate of excitatory neurons to L2/3 pyramidal cells plotted against the presynaptic cell’s distance from the pia for the indicated Cre lines. **(C)** Heatmap of translaminar connection rate for all excitatory Cre lines. White triangles represent locations of recorded L2/3 pyramidal neurons. Black regions correspond to two-dimensional bins with fewer than 10 connections probed. **(D)** Connection rate plotted against the absolute value of the lateral distance between pre- and postsynaptic neurons for neurons in L4 or L5. Shading indicates 95% confidence intervals.

Excitatory L6 corticothalamic (CT) neurons display limited axonal projections to superficial cortical layers (Kim et al., 2014; Bortone et al., 2014) and stimulation of L6 by glutamate uncaging suggests excitatory synaptic input from L6 to L2/3 is weak (Xu et al., 2016). We assayed connectivity from L6 CT neurons to L2/3 pyramidal neurons using two-photon stimulation of ChrimsonR-expressing cells in the Ntsr1-Cre line (Vélez-Fort et al., 2014). Consistent with the weak axonal projections (Kim et al., 2014; Bortone et al., 2014), we found a single excitatory connection from 1142 connections tested (Figure 3B). In further support of relatively weak projections from L6 compared to layers 4 and 5, responses to one-photon stimulation were dramatically smaller in Ntsr1 experiments than Rorb or Tlx3 experiments (Ntsr1: 0.121 ± 0.174 mV; Rorb: 9.09 ± 7.19 mV; Tlx3: 7.98 ± 5.62 mV) (Figure 3 - Supplement 1).

Figure 3C summarizes connectivity data obtained from all excitatory Cre lines used in this study - i.e., the probability of excitatory input to L2/3 neurons as a function of the (1) vertical distance from the pia and (2) horizontal distance between each putative synaptically-coupled pair of neurons. The spatial distribution of connected neurons was similar when connectivity was determined by any one of three methods: 1) visual assessment of photostimulus responses for a reliable EPSP (Figure 3B-D), 2) classification by a support vector machine (SVM) trained on a subset of the data, or 3) measurement of the peak stimulus-evoked voltage response relative to background variance (Figure 3 - Supplement 2). To examine to influence of lateral distance on the rate of translaminar excitatory connections, we measured connection rate against the absolute value of the lateral offset between pairs of neurons for stimulated presynaptic cells in L4 and L5 (Figure 3D). Connection rates from both L4 and L5 neurons displayed a significant dependence on the horizontal distance between pairs (L4: p=9.1e-10, L5: p=0.011, chi square test), although the effect was more dramatic for neurons in L4 (Chi square test statistics: L4 = 50.9, L5 = 14.8). More generally, it is evident from Figures 3C,D that excitatory cells with the highest connection rate to L2/3 pyramidal neurons are found in L4 with less than 200 μm of lateral offset between cells. By comparison, the more gradual decline in excitatory connection rate from L5 may suggest that L5 cells from a broader range of retinotopic locations project to L2/3 pyramidal cells.

### Translaminar and within layer inhibition by Sst neurons

How, if at all, does the pattern of vertical inhibitory input to L2/3 pyramids resemble that of vertical excitatory input to the same cells? The diverse axonal projection patterns and translaminar synaptic inhibition from L5 Sst neurons to L4 excitatory neurons (Kapfer et al., 2007; Silberberg and Markram, 2007; Nigro et al., 2018; Gouwens et al., 2019; Naka et al., 2019) inspired us to measure Sst-mediated inhibition to L2/3 pyramidal neurons and compare connection rates and strengths between intralaminar and translaminar inhibition. An example experiment utilizing AAV-mediated expression of soma-targeted-ChrimsonR shows a high rate of connectivity between Sst neurons in L2/3 and nearby pyramidal cells (Figure 4A). Consistent with previous reports (Fino and Yuste, 2011), we found a high rate of within layer connectivity that had a steep dependence on the lateral distance between cells using both transgenic and viral ChrimsonR expression (Figure 4B,C) (L2/3 within 100 μm lateral distance: Ai167: 20/43, 46.5%; AAV: 24/63, 38.1%; p=0.43, Fisher’s Exact). Nota l, we also o served translaminar inhi ition from Sst neurons in layers 4 and 5 to L2/3 pyramidal cells (Figure 4B,C), although the connection rate was only one-third that of intralaminar connections (L2/3 within 100 μm lateral distance: 44/106, 41.5%; L4 and L5 within 100 μm lateral distance: 28/200, 14%; p=1.8e-7, Fisher’s Exact). hus, inhi ition of L2 ramidal neurons Sst-interneurons originates primarily, but not exclusively from nearby Sst-neurons in the same layer.

**Figure 4.**
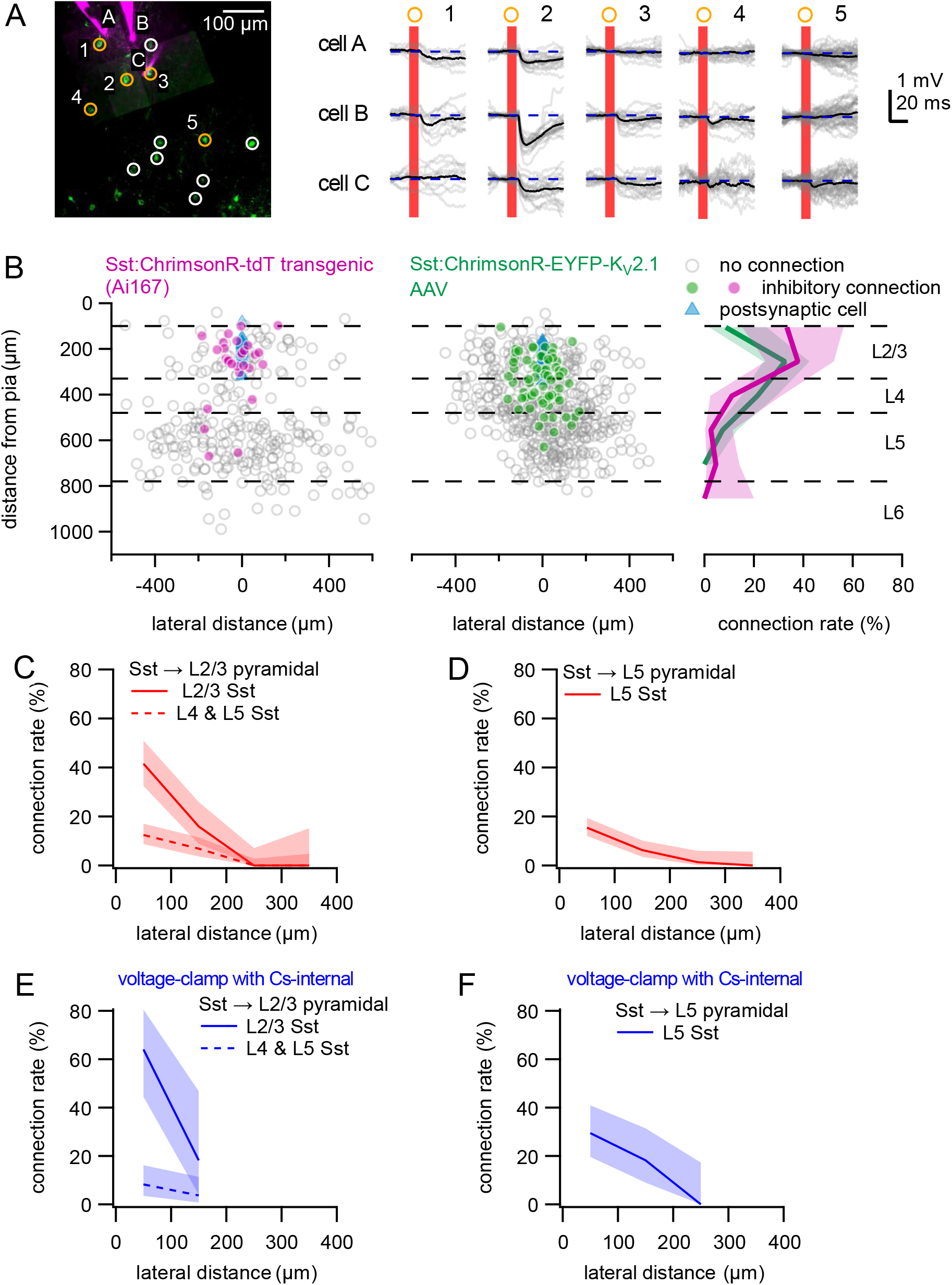
Measurement of inhibitory connectivity from L5 and L2/3 Sst interneurons to L5 and L2/3 pyramidal cells. **(A)** Example experiment measuring connectivity from Sst neurons to three L2/3 pyramidal neurons. *Left*: Flattened z-stack of recorded L2/3 pyramidal neurons filled with Alexa 594 (magenta) and EYFP signal from soma-targeted opsin-expressing neurons (green). Yellow circles indicate photostimulated cells with responses plotted in right panels. Responses following stimulation of cells in white circles are not shown. *Right*: Responses of cells to photostimulation of putative presynaptic partners. Postsynaptic cells were depolarized to −55 mV with automated bias current to increase the driving force of inhibitory currents. Photostimulus occurred during the red bar, black lines are an average of individual sweeps (grey). **(B)** Summary maps for all experiments with Sst:Ai167 (*left*) and Sst:AAV-injected (*center*) mice. Locations of neurons are plotted according to their distance from the pia and the lateral distance between putative pre- and postsynaptic cells with the following color scheme: *blue:* postsynaptic (recorded) L2/3 pyramidal cells *grey*: photostimulated neurons that did not evoke a synaptic response in the recorded cell. *magenta/green*: Stimulated cells that produced an IPSP using Ai167 or AAV-mediated opsin expression. Dashed lines represent approximate layer boundaries. Positive lateral distances correspond to presynaptic cells posterior to the postsynaptic cell. *Right:* The connection rate of Sst neurons to L2/3 pyramidal cells plotted against the presynaptic cell’s distance from the pia. Measurement of connection rate versus pia distance was limited to cells <200 μm lateral distance. **(C)** Connection rate versus lateral distance between Sst and L2/3 pyramidal neurons for Sst neurons within L2/3 (solid line) or in layers 4-5 (dashed line). **(D)** Connection rate versus lateral distance between cells for L5 Sst to L5 pyramidal neurons. **(E,F)** Connection rate as in panels C,D measured using Cs-based internal solution and voltage-clamp recordings. Shading in panels B-F represents 95% confidence intervals.

The high rate of connectivity observed in L2/3 resembles the high connection probability previously described for Sst-interneuron connectivity in frontal cortex (Fino and Yuste, 2011; Karnani et al., 2014). However, other studies have suggested that Sst-interneuron connectivity within L5 of somatosensory cortex is more selective (Nigro et al., 2018; Naka et al., 2019). To directly compare intralaminar Sst-interneuron to pyramidal neuron inhibition between L2/3 and L5 of mature mouse V1, we measured L5 Sst-interneuron to L5 pyramidal neuron connectivity, again using both Ai167 and AAV-mediated opsin expression (Figure 4D). We observed a dramatically lower rate of intralaminar Sst-interneuron to pyramidal neuron connectivity in L5 than in L2/3. (Figure 4C,D; L5 within 100 μm lateral distance: L2/3: 44/106, 41.5%; L5: 56/368, 15.2%, p=3.2e-8 Fisher’s Exact). Intralaminar inhibition from Sst-interneurons to pyramidal neurons in L5 was similar using either Ai167 or AAV-mediated ChrimsonR expression (within 100 μm of lateral distance, Ai167: 25/147, 17.0%; AAV: 31/221, 14.0%, p=0.55 Fisher’s Exact test). Layer-dependent differences in inhibitory connectivity were significant when connectivity was determined by either visual annotation of photoresponses, machine categorization, or measurement of maximum voltage responses relative to background variance (Figure 4 - Supplement 1).

The axons of many Sst neurons are thought to innervate the distal dendrites of both L5 and L2/3 neurons in L1. We wondered if the differences in intralaminar connection rates could be explained by the longer electrotonic length of L5 pyramidal neuron dendrites. Measurement of synaptic responses is inherently biased by somatic recordings in either current-clamp or voltage-clamp configurations (Spruston et al., 1993; Williams and Mitchell, 2008); however, it has been suggested voltage-clamp recordings may be more suitable for detection of distally-located or weak synapses (Barth et al., 2016). To determine if the observed difference in intralaminar connectivity was influenced by use of current-clamp recordings, we made additional voltage-clamp recordings from both L2/3 and L5 pyramidal neurons with a Cs-based internal solution including the sodium-channel blocker QX-314 (using Sst:Ai167 mice). As with current clamp experiments, intralaminar Sst-interneuron to pyramidal neuron connectivity was significantly higher in L2/3 than in L5 (Figure 4E,F, Figure 4 - Supplement 2; within 100 μm lateral, L2/3: 16/25, 64%; L5: 20/68, 29.4%, p=0.004). Our results using both recording strategies support differences in intralaminar Sst- to pyramidal-neuron connectivity between L2/3 and L5.

### Distributions of synaptic connection strengths

As shown in example experiments (Figure 2A, Figure 3A, Figure 4A), amplitudes of synaptic responses vary dramatically. We examined the probability density function of synaptic connection strengths measured in L2/3 pyramidal neurons for both excitatory connections (combining data from Rorb, Tlx3, Scnn1a and Ntsr1 Cre lines) and inhibitory connections from Sst-interneurons (Figure 5A,B). IPSP amplitudes are reported as their absolute value to facilitate comparison of amplitude distributions and distance correlations between excitatory and inhibitory responses. Distributions of both EPSP and IPSP amplitudes were better described by a lognormal function than a normal function (Figure 5A,B, Table 1). One implication of a lognormal distribution is that while strong connections are rare, they make a highly disproportionate contribution to the total synaptic strength received by a neuron. If we consider the cumulative strength of all translaminar excitatory connections measured onto L2/3 pyramidal neurons, 50% of this synaptic drive was generated by 13% of connections (corresponding to EPSP amplitudes > 0.82 mV). Inhibitory input from Sst neurons was only somewhat less skewed with 50% of synaptic drive coming from the strongest 19% of all connections.

**Figure 5.**
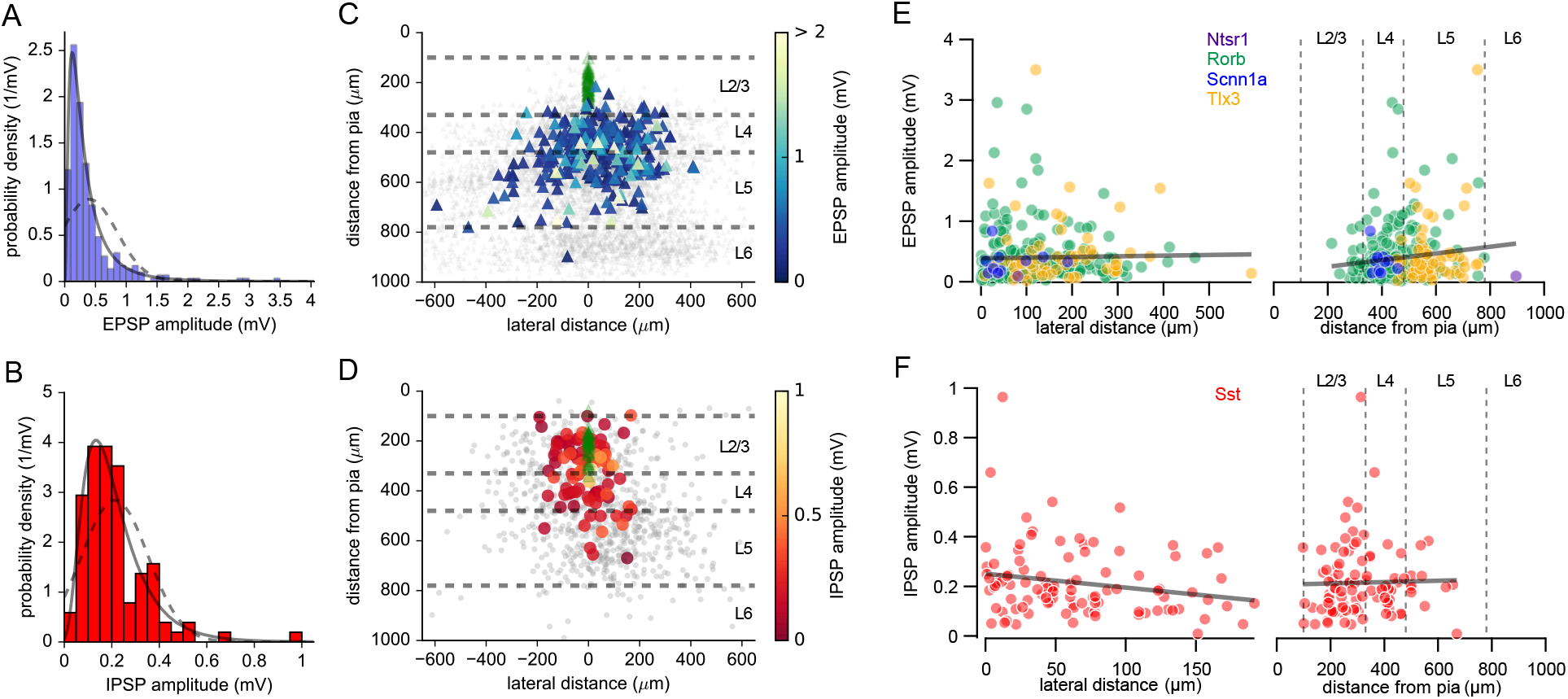
PSP amplitude distributions and distance dependence. **(A)** Probability density function of EPSP amplitudes measured from Ntsr1, Rorb, Scnn1a and Tlx3 Cre lines (blue). The distribution of EPSP amplitudes was fit by both normal (dashed line) and lognormal (solid line) distributions (see also Table 1). **(B)** Probability density function of IPSP amplitudes measured from Sst neurons as described for panel A. **(C)** Heatmap of EPSP amplitudes plotted according to lateral distance between pre- and postsynaptic cells and the presynaptic neuron’s distance from the pia. Dashed lines represent approximate layer boundaries. Locations of postsynaptic cells are plotted in green. **(D)** Heatmap of IPSP amplitudes as described for panel B. **(E)** EPSP amplitude plotted against lateral distance and distance from the pia for excitatory inputs. Markers are color coded by Cre line. Grey line represents linear regression of EPSP amplitude against corresponding distance measure. **(F)** IPSP amplitudes from Sst neurons plotted as in panel E.

**Table 1.** Characterization of EPSP and IPSP amplitude distributions. K-S stat. and K-S p-value refer to the Kolmogorov-Smirnov statistic and associated p-value obtained by comparing the measured distribution to the distribution obtained by fitting data with either a normal or lognormal distribution.

| *fit function* | *normal* |  |  |  | *lognormal* |  |  |  |  |
| --- | --- | --- | --- | --- | --- | --- | --- | --- | --- |
| fit parameter and goodness | mean | std. dev. | K-S stat. | K-S p-value | shape | loc | scale | K-S stat | K-S p-value |
| **excitatory to L2/3** | 0.40 | 0.45 | 0.22 | 3.09E-13 | 0.87 | -3.12E-03 | 0.27 | 0.06 | 0.32 |
| **Sst to L2/3** | 0.21 | 0.14 | 0.14 | 0.03 | 0.50 | -0.04 | 0.22 | 0.05 | 0.98 |

We also examined synaptic connection strengths as a function of the lateral distance between connected cells and the presynaptic neuron’s distance from the pia (Figure 5C-F). Overall, we find little to no correlation between PSP amplitudes and these distance measures (assessed by either parametric or non-parametric methods, Table 2). EPSP amplitudes measured in response to photostimulation of L5 excitatory neurons were on average larger, but not statistically different from those in L4 (L4: 0.38 ± 0.41 mV, L5: 0.43 ± 0.48 mV, Mann Whitney p=0.30). Similarly, the strengths of within layer and translaminar connections from Sst neurons were not statistically different (L2/3: 0.23 ± 0.16 mV, L4-5: 0.20 ± 0.11 mV, Mann Whitney p=0.35). Therefore, while vertical inhibition from deeper layer Sst neurons may be less common than within layer lateral inhibition, the influence of an individual Sst neuron on a L2/3 pyramidal cell does not appear to be determined by the layer it resides in.

**Table 2.** Distance dependence of EPSP and IPSP amplitudes. The strength and significance of correlations between PSP amplitudes and lateral distance or the presynaptic neuron’s distance from the pia. An α of 0.0125 was used to assess statistical significance (Bonferroni correction for 4 comparisons).

| tested correlation/strength & p-values | Pearson's R | Pearson's p | Spearman's rho | Spearman's p |
| --- | --- | --- | --- | --- |
| **excitatory amp. vs. lateral distance** | 0.03 | 0.65 | 0.16 | 0.01 |
| **excitatory amp. vs. distance from pia** | 0.13 | 0.03 | 0.06 | 0.35 |
| **Sst amp. vs. lateral distance** | -0.22 | 0.03 | -0.24 | 0.02 |
| **Sst amp. vs. distance from pia** | -0.01 | 0.89 | 0.05 | 0.65 |

### Locations and strengths of connections sharing post- or presynaptic neurons

Two-photon stimulation allowed us to probe up to 75 putative presynaptic partners of a recorded postsynaptic cell - many more than would have been feasible by patch clamping alone - and we observed 96 examples in which multiple presynaptic neurons (up to 12 connections per cell) converged onto individual L2/3 pyramidal cells (Figure 6A, also see examples in Figure 2A and Figure 4A). To examine the spatial dependence of convergence, we considered every unique pair of putative presynaptic neurons that were photostimulated while recording from a given postsynaptic cell. If both cells produced a synaptic response in the recorded cell, the pair was considered to be converging. For all pairs, we calculated the average lateral offset between each presynaptic cell and the postsynaptic target (Figure 6B) and the lateral offset between the two presynaptic cells (Figure 6C). Similar to overall connectivity rates, the rate of convergence was dependent on the lateral distance between putative presynaptic pairs and the postsynaptic target (Figure 6B). While this reinforces the observation that connectivity is most common between vertically aligned cells, we also observed instances in which presynaptic neurons separated by hundreds of microns in the lateral dimension converged onto a common L2/3 pyramidal cell (Figure 6C).

**Figure 6.**
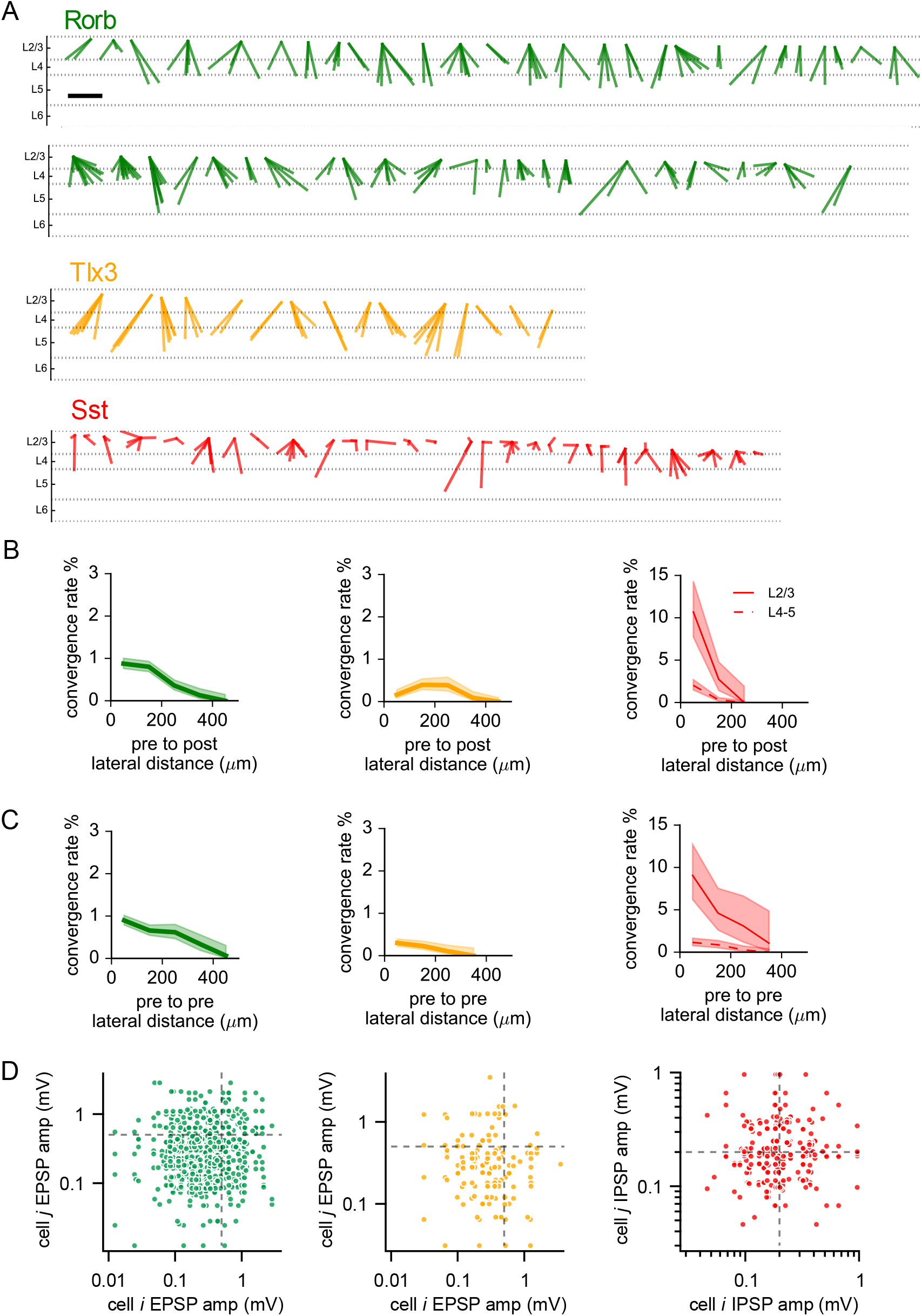
Convergence of synaptic inputs. **(A)** All identified instances of convergent inputs to L2/3 pyramidal neurons from Rorb (green), Tlx3 (orange) or Sst (red) labeled neurons. The same colors are used to indicate Cre lines throughout the figure. Lines are drawn from identified presynaptic partners to a common point representing the L2/3 pyramidal neuron. Cases of convergence are ordered according to the postsynaptic cell’s distance from the pia. Scale bar = 300 μm. **(B)** The rate of convergence plotted against the average lateral distance between each presynaptic neuron and the postsynaptic neuron for each Cre line. Sst neurons were split according to layer. **(C)** The rate of convergence plotted against lateral distance between pairs of presynaptic neurons. Shading represents 95% confidence intervals. **(D)** Scatter plots of incoming connections strengths for all pairs of cells sharing a postsynaptic partner for each Cre line.

Measurements of recurrent intralaminar connectivity between pyramidal neurons show that strong connections are frequently observed between bidirectionally connected pairs, and the strengths of connections sharing a presynaptic or postsynaptic neuron are correlated, suggesting that strong connections are clustered to a subset of cells (Song et al., 2005; Cossell et al., 2015). To determine if this was true of connections formed on L2/3 pyramidal neurons from Rorb, Tlx3, or Sst cells, we analyzed the relationship between the strengths of connections converging onto L2/3 pyramidal neurons (Figure 6D). The strength of connections sharing a postsynaptic target were not correlated for any of the three Cre lines. (Rorb: R=0.036, p=0.49, Tlx3: R=−0.0 2, =0. 0, Sst =0.0, =0.69, pearson’s measured in log-space). Therefore, while the overall distribution of connection amplitudes suggests a disproportionate influence of rare and strong synaptic connections (Figure 5), we do not find evidence that strong synaptic connections (of the classes examined here) are biased to particular L2/3 pyramidal neurons.

After finding that excitatory and inhibitory inputs of widely varying amplitudes converge across hundreds of microns to single L2/3 pyramidal cells, we next asked if the spatial distribution of divergence from single presynaptic neurons to multiple L2/3 pyramidal neurons is similarly broad. While the number of simultaneous recordings limits the number of connections we may find from a given presynaptic cell to 4, the relatively large number of photostimulated presynaptic neurons permits us to test many instances of divergence. Figure 7A illustrates all instances of divergence to L2/3 pyramidal neurons observed when targeting presynaptic cells from the indicated Cre lines (also see examples in Figure 3A, 4A). For Rorb and Tlx3 excitatory neurons, the observed rates of divergence do not depend on the intersomatic distance between postsynaptic neurons (Figure 7B) (Rorb: <100 μm: 9/466, 100-200 μm 21, Fisher’s Exact: p=1.0; Tlx3: <100 μm: 3/315, 100-200 μm, 1 2 9, Fisher’s Exact: p=0.6). his demonstrates that the axons of single excitatory neurons form contacts onto L2/3 pyramidal cells hundreds of microns away from one another. In contrast, single L2/3 Sst neurons are more likely to contact pairs of L2/3 pyramidal cells that are near each other than pairs 100-200 μm apart (<100 μm: 12/43, 100-200 μm, Fisher’s Exact p=0.012). he relatively high, but steeply declining, rate of inhibitory divergence suggests that connectivity of L2/3 Sst neurons is less selective than translaminar excitatory input, but more spatially confined. Divergence from Sst neurons in deeper layers was uncommon (consistent with the lower connection rate) and the rate was similar when recorded cells were nearby or relatively far apart (<100 μm: 2/165, 100-200 μm 1, Fisher’s Exact p=0.43).

**Figure 7.**
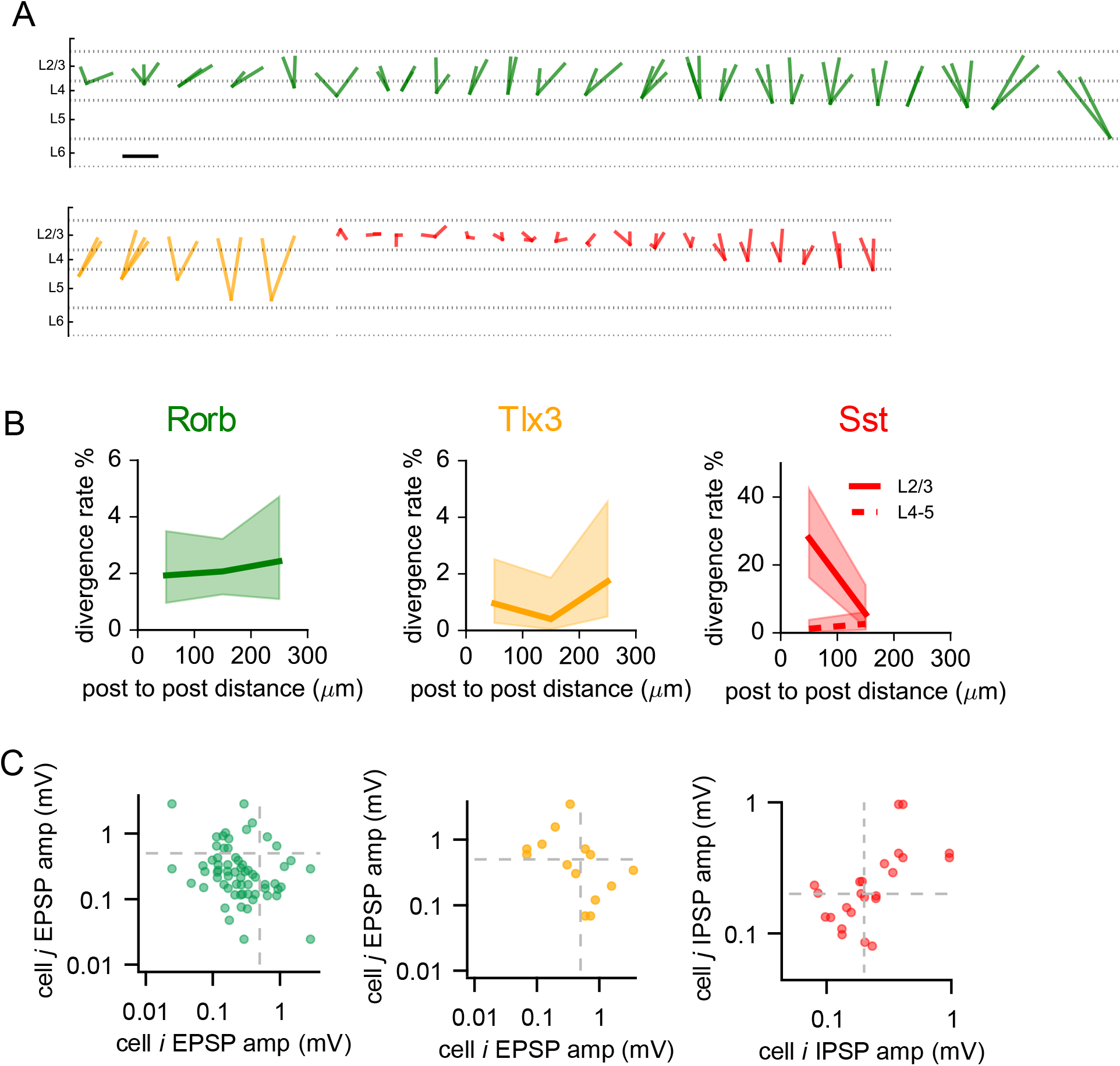
Divergence of synaptic outputs. **(A)** All identified instances of divergent output to multiple L2/3 pyramidal neurons from Rorb (green) Tlx3 (orange) and Sst (red) labeled neurons. The same colors are used to indicate Cre lines throughout the figure. Lines are drawn from presynaptic neurons to 2-4 points representing the connected L2/3 pyramidal cells. Cases of divergence are ordered according to the presynaptic cell’s distance from the pia. Scale bar = 300 μm. **(B)** The rate of divergence plotted against the intersomatic distance between pairs of postsynaptic neurons. Shading represents 95% confidence intervals. **(C)** Scatter plots of outgoing connections strengths for all pairs of cells sharing a presynaptic partner for each Cre line.

Finally, we examined the relationships between the strengths of divergent connections made by individual presynaptic cells onto L2/3 pyramidal cells (Figure 7C). We did not observe significant correlations when examining EPSP amplitudes from single Rorb or Tlx3 neurons onto multiple L2/3 pyramidal neurons (Rorb: R=-0.28, p=0.12; Tlx3: R=−0.46, p=0.30). We did however find a moderate positive correlation in the amplitudes of synaptic responses produced by diverging Sst-interneurons (Sst: R=0.62, p=0.03). In other words, an Sst neuron that forms a strong connection onto one L2/3 pyramidal neuron is likely to also form strong connections onto other L2/3 pyramidal neurons.

## Discussion

We have combined multicellular recording with two-photon optogenetic stimulation to characterize translaminar excitatory and inhibitory inputs to L2/3 pyramidal neurons in mature mouse V1. We find that ChrimsonR is a suitable opsin for photoactivation by rapid two-photon scanning in several Cre-line defined subclasses of neurons, though we did observe some differences in photosensitivity across Cre lines (Figure 1, Figure 1 - Supplement 1). Photosensitivity was higher for Sst neurons with AAV-meditated opsin expression than transgenic expression - though the same effect was not observed for pyramidal neurons. The reliability of light-evoked spiking has clear implications in interpretation of experiments using two-photon photostimulation to measure synaptic connectivity as connections will not be detected if the presynaptic cell is not firing. In the context of results presented here, we found the fewest connections when examining connectivity from Ntsr1 neurons (L6 CT cells) to L2/3 pyramidal neurons, in agreement with previous reports of little to no excitation arising from L6 neurons (Dantzker and Callaway, 2000; Xu et al., 2016). Ntsr1 provided the most reliable and precise generation of light-evoked APs (Figure 1, Figure 1 - Supplement 1), therefore false-negatives associated with two-photon stimulation are unlikely to explain this result.

Another potential source of false negatives (inherent to acute slice preparations and not unique to photostimulation) is the severing of neuronal processes. This subject has been explored in other studies and in this context, connection rates measured in vitro may be thought of as lower bounds for connection rates present in vivo (Stepanyants et al., 2009; Levy and Reyes, 2012). In this study, slices were cut at a slightly oblique angle to help preserve neuronal processes and we targeted cells for both patching and photostimulation that were >40 μm below the surface of the slice (Methods). We observed connections from excitatory neurons in L5 to L2/3 neurons with 300-600 μm of lateral offset (10 connections/489 probed, 2%) - corresponding to total intersomatic distances of 484-735 μm. Considering this, we believe the absence of connections observed over similar intersomatic distances (e.g. from Sst neurons with >200 μm of lateral offset or from Ntsr1 neurons) represents selectivity of the connections and is not strictly a result of acute slice preparation. However, we acknowledge that intersomatic distance is not equivalent to the distance an axon may travel to form a synaptic contact (e.g. synapses formed on the distal dendrites of a nearby postsynaptic cell), and some classes of connections may be more prone to cutting than others due to tortuous axon trajectories.

The sub-cellular location of synaptic contacts also relates to the potential to miss distally located (or otherwise weak) connections when measuring synaptic responses by somatic recordings. To this end, recording of postsynaptic responses in either voltage-or current-clamp each come with inherent advantages and disadvantages (Williams and Mitchell, 2008; Barth et al., 2016; Jiang et al., 2016). Although voltage-clamp recording is generally thought to increase the ability to detect small responses (especially when combined with pharmacology to isolate synaptic currents), it presents some challenges in the interpretation of results due to space-clamp errors (Spruston et al., 1993; Williams and Mitchell, 2008; Beaulieu-Laroche and Harnett, 2018). Additionally, use of ion channel blockers to improve space clamp limits the ability to characterize the intrinsic properties of recorded cells. In this study, the relatively high throughput of our approach enabled us to perform both current-clamp measurements of IPSPs using a physiological K-gluconate-based internal solution and voltage-clamp measurements of isolated inhibitory postsynaptic currents (IPSCs) using a Cs-based internal solution to compare Sst to pyramidal neuron connectivity in L2/3 and L5 (Figure 4, Figure 4 - Supplements 1 and 2). It was especially valuable to use both methods in this context given the axonal projection patterns of Martinotti-type Sst neurons to L1 and likelihood of distal synaptic contacts. Both datasets indicate a significantly higher rate of intralaminar Sst to pyramidal neuron connectivity in L2/3 than in L5 and the presence of translaminar inhibition from subgranular Sst neurons to L2/3 pyramidal cells.

There is a retinotopic map of visual space that is conserved across layers in V1 - arguing for strong vertical synaptic connections (Dräger, 1975; Mangini and Pearlman, 1980; Wagor et al., 1980; Schuett et al., 2002; Bonin et al., 2011). Rodent V1 lacks the smoothly varying orientation columns observed in higher mammals (Grinvald et al., 1986; Hübener et al., 1997; Ts’o et al., 1990), instead neurons with disparate orientation tuning are intermingled in a “salt and pepper” fashion (Ohki et al., 2005). However, it has also been reported that nearby cells display more similar tuning than those farther away from each other (Ringach et al., 2016) and that local biases in orientation tuning reveal a global map in orientation tuning that spans V1 and neighboring visual areas (Fahey et al., 2019). Synaptic connections with little lateral offset may act to preserve or stabilize functional maps, while connections between more horizontally-distributed neurons may promote their reorganization. The highest rate of excitatory neuron connectivity to L2/3 pyramidal neurons was observed from L4 neurons with 0-100 μm of lateral offset - a likely means of preserving the retinotopic map established by inputs from dLGN (Figure 3C,D). However, the falloff in connection rate with lateral offset was gradual for both L4 and L5 excitatory inputs to L2/3 pyramidal neurons (Figure 3C,D). Additionally, there was little to no correlation between intersomatic lateral offsets and EPSP amplitudes (Figure 5). These findings, along with our observation of both convergence and divergence across hundreds of microns of horizontal space (Figure 6,7), illustrates putative synaptic mechanisms by which translaminar excitatory connections can promote both the maintenance and redistribution of sensory information across layers.

We found that excitatory input from L5 IT neurons to L2/3 pyramidal neurons rivals excitation from L4. While we did not observe a significant difference in the average EPSP amplitudes generated by L4 or L5 excitatory neurons, the fraction of cells that produced a strong synaptic response (>1 mV) in L5 was nearly twice that of L4 (5.8% versus 10.8% of all connections in layers 4 and 5 respectively). The strength of connections from L5 IT cells to L2/3 pyramidal neurons is interesting in the context of *in vivo* studies demonstrating strong influence of brain state and active behavior on the activity of neurons in primary visual cortex (Niell and Stryker, 2010; Keller et al., 2012; Neske et al., 2019; Stringer et al., 2019). If L5 neurons are receiving long range input encoding behavioral variables from higher cortical areas, the strong (Figure 5) and broadly diverging (Figure 7) local projections from L5 IT neurons could provide a mechanism to distribute this information within V1 so that it can be integrated with projections providing sensory information (e.g. thalamic input or L4 to L2/3 connections).

Why were excitatory connections from L5 to L2/3 previously thought to be sparse or weak? The age range of animals used in this study is older than most. Connectivity in younger animals may be more specifically driven by sensory input. It is also possible that use of Tlx3-Cre (and Rorb-Cre) allowed us to more thoroughly identify and measure these somewhat rare connections that may be specific to IT-type L5 neurons, especially in visual cortex where L5 excitatory types are not as distinctly separated into sub-layers compared to somatosensory cortex (Groh et al., 2010; Kim et al., 2015). The ability to probe connectivity of genetically-defined subclasses of neurons with single cell resolution represents a major advantage of this approach over photostimulation via one-or two-photon glutamate uncaging. Measurement of local connectivity via two-photon optogenetics in future studies will benefit from further use of available Cre-driver lines to express opsin proteins in defined classes of neurons. Furthermore, there is the potential to test for differences in synaptic connectivity between transcriptomically-defined cell types that exist within Cre-line-defined-subclasses by following two-photon circuit characterization with *post hoc* measurement of gene expression in the very same neurons using multiplex fluorescence *in situ* hybridization (Nicovich et al., 2019).

This study illustrates that combining multi-cellular recording with multi-photon optogenetic stimulation is an effective means of characterizing synaptic connectivity between genetically-defined populations of neurons. On this topic, the Allen Institute recently released a freely available dataset - focused on intralaminar connections - describing the rate, strength, and short-term dynamics of synaptic connectivity measured by multi-cellular recording (https://portal.brain-map.org/explore/connectivity/synaptic-physiology). Future work utilizing a variety of complementary electrophysiological and optical methods to characterize local and long-range synaptic signaling between categories of neurons promises to further our understanding of cell types and their unique roles in circuit function and behavior.

## Methods

### Animals and stereotaxic injections

Mice were housed and sacrificed according to Institutional Animal Care and Use Committee approved protocols. Cre driver lines used in this study include Ntsr1-Cre_GN220, Rorb-IRES2-Cre, Scnn1a-Tg3-Cre, Sst-IRES, and Tlx3-Cre_PL56. Driver names were simplified as Ntsr1, Rorb, Scnn1a, Sst, and Tlx3 throughout the manuscript. Transgenic expression of ChrimsonR was achieved by crossing the indicated Cre line to the effector line Ai167(TIT2L-ChrimsonR-tdT-ICL-tTA2) (Daigle et al., 2018). Expression of soma-targeted-ChrimsonR was achieved by stereotaxic injection of AAV serotype 1 carrying ChrimsonR-EYFP fused to a Kv2.1 somatic lo alization motif (“fle - ChrimsonR-EYFP-kv”, Addgene plasmid #135319). Injections were made into right visual cortex of mice aged between P26-P53. Injection coordinates were 3.8 mm posterior from bregma and 3.0 mm lateral from midline. Two injections of 100 nL each were targeted 300 and 600 μm below the surface of the pia with the goal of transfecting cells in all cortical layers. Experiments were performed 21-28 days after injection.

### Slice preparation

Brain slices were prepared from young adult mice (P40-P80) of either sex. Mice were anesthetized with isoflurane then transcardially perfused with oxygenated, cold NMDG-slicing solution containing (in mM): 98 HCl, 96 N-methyl-d-glucamine (NMDG), 2.5 KCl, 25 D-Glucose, 25 NaHCO_3_, 17.5 4-(2-hydroxyethyl)-1-piperazineethanesulfonic acid (HEPES), 12 N-acetylcysteine, 10 MgSO_4_, 5 Na-L-Ascorbate, 3 Myo-inositol, 3 Na Pyruvate, 2 Thiourea, 1.25 NaH_2_PO_4_ H_2_0, 0.5 CaCl_2_ and 0.01 taurine. Acute arasagittal sli es (0 μm thi k) were ut with either a om resstome (re isionar Instruments) or a Leica VT1200S at an angle of 17° relative to the sagittal plane to preserve pyramidal cell apical dendrites (Seeman et al., 2018). Slices were stored at 34° C in NMDG-slicing solution for 10 minutes, after which slices were maintained at room temperature in holding solution containing (in mM): 94 NaCl, 25 D-Glucose, 25 NaHCO_3_, 14 HEPES, 12.3 N-acetylcysteine, 5 Na-L-Ascorbate, 3 Myo-inositol, 3 Na Pyruvate, 2.5 KCl, 2 CaCl_2_, 2 MgSO_4_, 2 Thiourea, 1.25 NaH_2_PO_4_ H_2_0, 0.01 Taurine.

### Electrophysiological recordings

At least 1 hour after slicing, slices were transferred to recording chambers with a constant perfusion of aCSF containing (in mM): 126 NaCl, 18 NaHCO_3_, 12.5 D-Glucose, 3 KCl, 2 CaCl_2_, 2 MgSO_4_, 1.25 NaH_2_PO_4_, 0.16 Na L-Ascorbate. pH was measured between 7.2 and 7.3 and osmolarity was between 290 and 300 mOsm. aCSF was bubbled with carbogen and maintained at 32-34° C with an inline heater. Slices were visualized under infrared illumination using either a 10x/0.2 NA or 40x/1.0 NA objective (Zeiss) and a digital camera (Hamamatsu; ORCA Flash4.0). Recordings were made from one to four electrode headstages that were amplified and low-pass filtered at 10 kHz using Multiclamp 700B amplifiers (Molecular Devices). Signals were digitized at 25-100 kHz using ITC-18 digitizers (Heka). Data was acquired using Multi-channel Igor Electrophysiology Suite (MIES; https://github.com/AllenInstitute/MIES), within Igor Pro (WaveMetrics).

Patch electrodes were pulled from thick-walled filamented borosilicate glass (Sutter Instruments) with a DMZ Zeitz-Puller. Tip resistance was 3–6 MΩ. Pipette recording solution contained (in mM): 130 K-gluconate, 10 HEPES, 0.3 ethylene glycol-bis(β-aminoethyl ether)-N,N,N’,N’-tetraacetic acid (EGTA), 3 KCl, 0.23 Na_2_GTP, 6.35 Na_2_Phosphocreatine, 3.4 Mg-ATP, 13.4 Biocytin. 25 μM of either Alexa-594 or Alexa-488 was added for experiments using ChrimsonR labeled with enhanced yellow fluorescent protein (EYFP) or tdTomato, respectively. Osmolarity was between 280-290 mOsm. pH was adjusted to 7.2-7.3 using KOH. For voltage-clamp experiments, K-gluconate was replaced by equimolar Cs-gluconate and 1 mM *N*-(2,6-Dimethylphenylcarbamoylmethyl)triethylammonium-Br (QX-314) was added. Reported voltages are uncorrected for a liquid junction potential of 9 mV between aCSF and the K-gluconate based internal and 10 mV between aCSF and the Cs-gluconate based internal.

For mapping experiments, one to four pyramidal neurons in either L2/3 or L5 were targeted for patching based on somatic appearance and depth from the surface of the slice (>40 μm). For most experiments, we measured synaptic responses evoked by one-photon stimulation of ChrimsonR-expressing neurons before conducting two-photon stimulation of individual cells (Figure 3 - Supplement 1). When measuring synaptic responses, automated bias current injection was used to maintain the membrane potential near −70 mV or −55 mV for mapping of excitatory or inhibitory inputs, respectively. Membrane potential was clamped to 0 mV for voltage clamp experiments measuring IPSCs.

Pyramidal cell identity was confirmed by examination of cell morphology in a two-photon z-stack acquired at the end of each experiment. Z-stacks were examined for dendritic spines and a prominent apical dendrite. To further assist in our classification of the postsynaptic cells, intrinsic properties were characterized by a family of 1 second square current injections from −130 pA to 250 to pA in 20 pA steps. Responses to current injection were measured from a baseline membrane potential of −70 mV (adjusted by steady bias current injection) (Table 3). Some cells required current injections >250 pA to evoke APs (again increased in 20 pA intervals). Intrinsic properties were measured as described by (Gouwens et al., 2019) with the exception that the smallest hyperpolarizing current injection (−10 pA) was not used in our estimation of the membrane time constant. Exponential fits to the weakest hyperpolarizing current injection were often inaccurate due to background synaptic activity present in our experiments. The average firing rate and adaptation index were measured from current injections 40 pA above rheobase. A small number of cells initially identified as pyramidal cells displayed electrophysiological characteristics typical of interneurons (high input resistance, narrow AP width, and/or small upstroke:downstroke ratio) and were not included in analyses of synaptic connectivity to pyramidal neurons.

**Table 3.** Intrinsic physiological properties of recorded neurons. Measured intrinsic values for L2/3 pyramidal neurons and L5 pyramidal neurons included in analyses of synaptic connectivity. Data are presented as mean ± standard deviation.

| cell class/measurement | L2/3 pyramidal | L5 pyramidal |
| --- | --- | --- |
| $V_{rest}$ (mV) | $-72.6 \pm 7.2$ | $-61.7 \pm 12.2$ |
| $R_i$ (MΩ) | $86.2 \pm 26.8$ | $103.1 \pm 39.6$ |
| membrane time constant (ms) | $14.2 \pm 4.5$ | $14.4 \pm 3.8$ |
| voltage sag fraction (0-1) | $.034 \pm .027$ | $0.2 \pm 0.13$ |
| AP width (ms) | $1.35 \pm 0.29$ | $1.21 \pm 0.23$ |
| upstroke:downstroke ratio | $4.35 \pm 0.66$ | $3.55 \pm 0.67$ |
| AP height (mV) | $89.4 \pm 10.5$ | $84.1 \pm 12.5$ |
| rheobase (pA) | $173 \pm 65$ | $144 \pm 61$ |
| F-I curve slope (spikes/s/pA) | $0.13 \pm 0.05$ | $0.16 \pm 0.62$ |
| avg. firing rate (spikes/s) | $7.8 \pm 2.3$ | $21.9 \pm 46.0$ |
| adaptation index | $0.09 \pm 0.07$ | $0.09 \pm 0.11$ |

### Photostimulation

Experiments were performed using Prairie Ultima two-photon laser scanning microscopes (Bruker Corp). A tunable pulsed Ti:Sapphire laser (Chameleon Ultra, Coherent) was used for imaging and a fixed wavelength (1070 nm) pulsed laser (Fidelity 2, Coherent) was used for photostimulation. Locations of putative presynaptic neurons were manually identified from two-photon reference images acquired in PrairieView software. The experimenter placed a target at the center of the soma, around which a spiral path of 10 μm diameter was generated. Stimulus duration was 10 ms and each cell was stimulated at least 20 times at 85 mW for all Cre-line opsin combinations except for Sst-Cre neurons transfected with AAV. Experiments using this Cre line and expression method were stimulated at 35 mW due to higher photosensivity of opsin expressing neurons (Figure 1). Photostimulation and electrophysiology recordings were synchronized via a TTL trigger generated by MIES acquisition software. The voltage output from the Prairie system to the Pockels cell was recorded in MIES for alignment of electrophysiological recordings to the photostimulus.

For experiments characterizing light-evoked generation of APs, loose-seal recordings (20-0 MΩ seal resistance) were made from opsin-expressing neurons 40-100 μm below the surface of the slice. Opsin-expressing cells were identified by brief reflected light illumination via an LED. Prior to two-photon stimulation, we confirmed our ability to detect light-evoked APs by applying a 1 ms one-photon stimulation to the cell via the 40x objective (1.3-1.5 mW, 590 nm). The minimum power required to drive reliable two-photon-evoked spiking was determined as the stimulus intensity that generated APs on 10 out of 10 trials.

To characterize the lateral resolution of our photostimulus, we delivered stimuli in a radial grid pattern containing seven spokes with stimuli 10, 20, and 30 μm away from the center of the recorded cell. To measure the axial resolution of our photostimulus, we offset the focus of the objective above and below the recorded cell in 10 μm increments until we observed spiking on 0 out of 10 trials. Spatial resolution was characterized using the same photostimulation powers used for mapping.

### Analysis of connectivity

The individual and averaged responses to photostimulation were quantitatively- and visually-assessed for evidence of a synaptic response. The features in Table 4 were measured in a 0 ms “post-stimulus” window eginning at the start of hotostimulation and a 0 ms “pre-stimulus” window 0 to 20 ms efore hotostimulation - when no evoked responses are expected (Figure 5 - Supplement 1). The standard deviation of the of the average response and the deconvolved response were measure in a 20 ms window immediately before the onset of the photostimulus. The features were used to train a SVM on human annotation of the presence or absence of a synaptic response (using the Sci-Kit Learn package in Python http://scikit-learn.org/stable/). Experiments examining excitatory or inhibitory responses were considered separately. The excitatory classifier was trained on 2363 randomly selected probed connections and tested against 3545 withheld probed connections. The classifier achieved 99% overall accuracy in classifying the test dataset (Figure 3 - Supplement 1). The inhibitory classifier was trained on 1182 examples and tested on 789 withheld examples and achieved 97% overall accuracy (Figure 4 - Supplement 1). False negatives missed by the classifier were typically reliable but low amplitude or slowly rising voltage responses, whereas false positives were often caused by large changes in membrane potential only present on a small number of trials. To determine if differences in human versus machine classification impacted our measurements of connectivity, we ran the classifiers on the full datasets and examined the spatial distribution of connection probabilities (Figure 3 - Supplement 1 and Figure 4 - Supplement 1). The classifications provided by the support vector machine displayed similar patterns of connectivity with regard to cortical layer of the presynaptic cell, Cre line and lateral offset between cells.

**Table 4.** Features measured for all photostimulus responses.

| Features from average response | Features from deconvolved average response | Features from individual deconvolved responses |
| --- | --- | --- |
| Maximum amplitude of response (peak) | Peak of deconvolved average (deconvolved peak) | average number of times each trace exceeded 3x deconvolved SD |
| Time of peak | Time of deconvolved peak | SD of number of crossings per sweep |
| Standard deviation (SD) at time of peak | SD of deconvolved trace in baseline window (deconvolved SD) |  |
| SD in baseline window | number of times the deconvolved trace exceeded 3x deconvolved SD |  |
|  | number of times the deconvolved trace exceeded 5x deconvolved SD |  |

As an alternative measure of connectivity, we generated post-stimulus and pre-stimulus z-scores for each photoresponse by dividing the peak amplitude of the average response by the standard deviation of the average response measured in a 20 ms baseline period (Figure 5 - Supplement 1). We then set a threshold equal to the 99th percentile of all pre-stimulus z-scores. If post-stimulus z-scores exceeded that threshold, corresponding cells were classified as connected. Discrepancies between manual annotation and this single measure were slightly higher compared to the machine classifier. Despite that, the patterns of connectivity examined by of presynaptic layer of origin, lateral offset or Cre line were similar for all 3 means of connection classification (Figure 3 - Supplement 1, Figure 4 - Supplement 1 and Supplement 2).

Stage coordinates of the photostimulated neurons were extracted from the position of the motorized stage and the targeted locations of the photostimulation galvos. Photostimulus locations were then offset to the position of the recorded neuron and rotated according to the orientation of the slice to provide vertical (along the pia to white matter axis) and lateral offsets between the pre- and postsynaptic neurons of every connection probed. The distance of the postsynaptic neuron from the pia was measured from overlaid transmitted light and fluorescence images captured after the neuron was filled with fluorescent dye. This postsynaptic distance from the pia was added to the vertical offset between probed cells to calculate the distance of the presynaptic neuron from the pia. This does not account for curvature of the pial surface, which was minimal for V1 across the lateral distances considered here. Representative layer boundaries used in this study are L1: 0-100 μm, L2/3: 100-330 μm, L4: 330-480 μm, L5: 480-780 μm, L6: 780-1000 μm. These boundaries were informed by the Allen Reference Atlas and the positions of presynaptic neurons assayed using the layer specific excitatory Cre lines in this study (see Figure 3). To measure connection rate as a function of the presynaptic neuron’s distance from the pia and lateral offset, we used 150 μm and 100μm bins respectively. Adjustment to bin sizes did not influence findings. We calculated 95% Jeffreys Bayesian confidence intervals using the number of connections found and probed (presented as shading throughout the manuscript). We used Fisher’s Exact tests when statistically comparing connection rates between two groups. Chi square tests were used to test the influence of distance on connectivity and to compare between more than two groups.

To measure convergence rate, we considered every unique pair of putative presynaptic neurons probed while recording from a given postsynaptic cell. If both neurons generated a synaptic response upon photostimulation, the pair was scored as one occurrence of converging presynaptic inputs. The distances between every potential converging pair were calculated to generate the convergence rate versus distance plots in Figure 6B,C. Pre to post distance was calculated as the average lateral offset between each presynaptic neuron and the common postsynaptic target. Pre to pre distance was calculated as the total lateral offset between the two presynaptic cells. Similarly, to measure divergence we considered every unique pair of simultaneously recorded postsynaptic neurons and identified pairs of cells that both received input from a common presynaptic partner as instances of divergence. The total distance between L2/3 pyramidal neurons was calculated to generate the plots in Figure 7B.

When measuring PSP amplitude correlations (Figure 6D and 7C), assignment of a connection strength as *i* or *j* would be arbitrary and potentially influence the measurement of Pearson’s R. Therefore, each pair of connections was lotted twice with a given connection amplitude assigned once as *i* and once as *j* and an *R* score was calculated from these values. To determine the probability of obtaining the measured *R* score (*p-*value), we used the number of unique pairs in the data (one half of the number of pairs plotted). A similar approach was taken to examine recurrent connectivity between L5 pyramidal neurons (Song et al., 2005).

### Analysis of PSP amplitudes

Voltage recordings were excluded from measures of connection strength if the voltage preceding the photostimulus differed from the target voltage by >5 mV or if the voltage changed by >1 mV in a 50 ms window preceding the photostimulus. The latter was done to avoid the influence of rare, large amplitude spontaneous events. Individual voltage recordings were baseline adjusted by subtracting the median membrane potential in a 20 ms window before photostimulation. An average PSP was generated from the baselined sweeps. The number of two-photon-evoked presynaptic spikes and their timing varies across photostimulated neurons (Figure 1C and Figure 1- Supplement 1). Therefore, to identify the timing of the first PSP and measure its amplitude, we used an exponential deconvolution of the average photoresponse to emphasize the timing of underlying synaptic events (equation 1) (Richardson and Silberberg, 2008). τ was set to 20 ms.

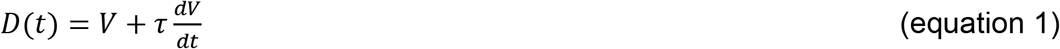

We identified peaks in the deconvolved trace that occurred 0-50 ms after the start of photostimulation. We set the threshold for peak detection to equal 5 times the standard deviation of the deconvolved trace prior to photostimulation (measured in a 20 ms pre-stimulus window). If a single threshold crossing was found, we measured the peak of the average voltage response in a time window from the first threshold crossing to 20 ms later. If multiple peaks were found, the window was refined to start at the first threshold crossing, and end at the time of the inter-peak-minimum.

The probability density of functions of EPSP and IPSP amplitudes were compared against normal and lognormal distributions using the scipy.stats Python package. Measured amplitudes were fit with normal and lognormal distributions. Resulting fit parameters are presented in Table 1, and corresponding distributions are plotted in Figure 5A,B. The goodness of each fit was determined by comparing the fit to the measured distribution using a Kolmogorov-Smirnov test.

## Acknowledgements

We thank the Allen Institute founder, Paul G. Allen, for his vision, guidance, and support. We are grateful for the assistance of the Allen Institute for Brain Science Neurosurgery and Behavior, Tissue Processing, and Transgenic Colony Management teams. We thank Lindsey Glickfeld, Alex Hoggarth, Brian Kalmbach, Scott Owen, and Stephanie Seeman for feedback on the manuscript.

## Supplemental Figures

**Figure 1 - Supplement 1.**
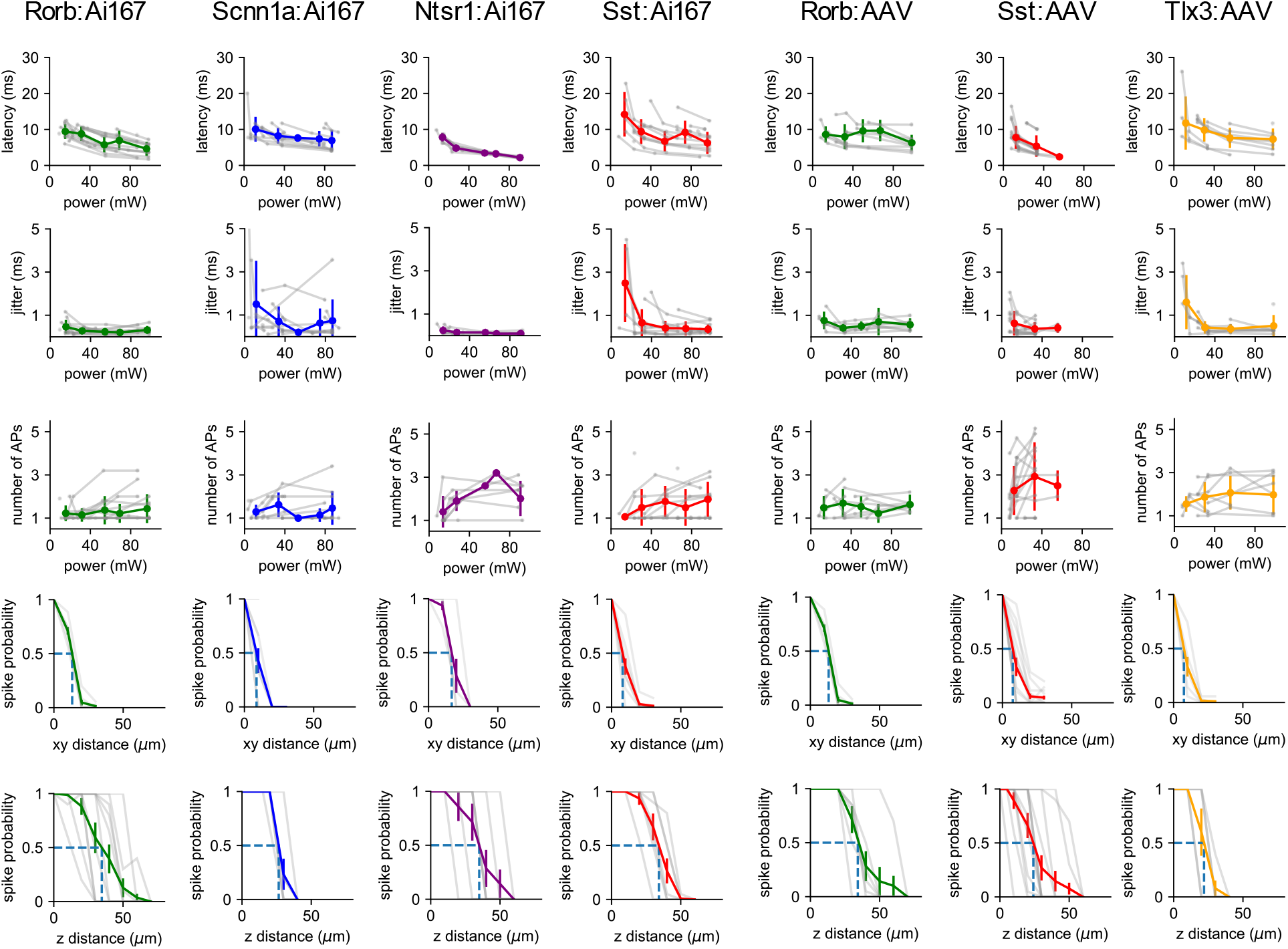
Characterization of spiking evoked by two-photon stimulation. Columns represent data collected from the indicated Cre line either crossed to the Ai167 effector line (ChrimsonR-tdTomato expressing) or injected with AAV carrying FLEX’ed soma-targeted-ChrimsonR (ChrimsonR-EYFP-Kv2.1). Each row represents the indicated measurement with data plotted for individual neurons (*grey*) and the average values measured for each Cre line/effector combination (*color*). For measurements of latency, jitter, and the number of APs versus power, average values were calculated by placing single cell measurements into 20 mW bins. Spatial resolution was characterized using the same photostimulation powers used for mapping. This power was 85 mW for all experiments except those using Sst-Cre with AAV injection. These cells exhibited greater photosensitivity (Figure 1B) and were therefore stimulated at 35 mW.

**Figure 3 - Supplement 1.**
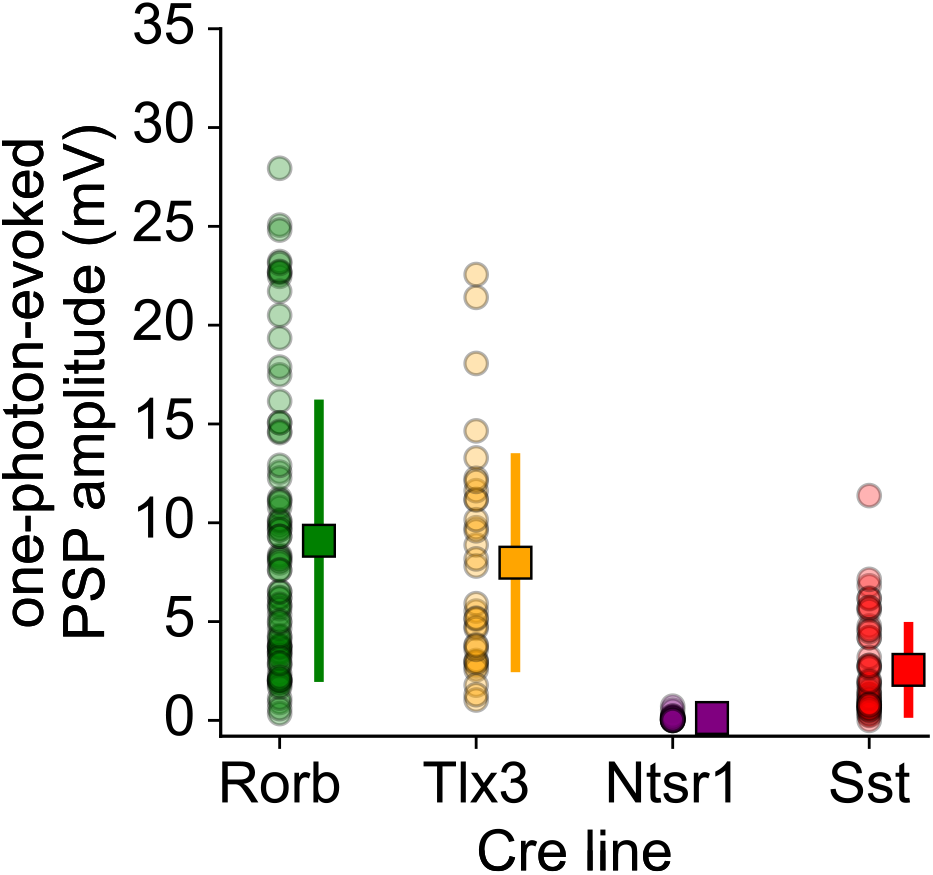
One-photon evoked synaptic responses. Amplitudes of postsynaptic responses measured in L2/3 pyramidal cells, evoked by one-photon stimulation of ChrimsonR-expressing neurons for experiments using the indicated Cre lines. Amplitudes were measured in response to a single 1 ms stimulus (590 nm) via a 40x objective centered over the cell. Data were collected from the same cells used for two-photon mapping. Circles represent values for individual neurons. Squares indicate mean ± standard deviation. Error bars for Ntsr1 are smaller than the marker. In Rorb-Cre experiments, 4 out of 82 recorded L2/3 pyramidal neurons generated an AP in response to summated synaptic inputs driven by one-photon stimulation. For these cells, the one-photon-evoked response was measured as the difference between AP threshold and the resting membrane potential. This indirect one-photon-evoked spiking was not observed in L2/3 pyramidal cells in experiments using Tlx3-Cre (33 recorded neurons) or Ntsr1-Cre (23 recorded neurons).

**Figure 3 - Supplement 2.**
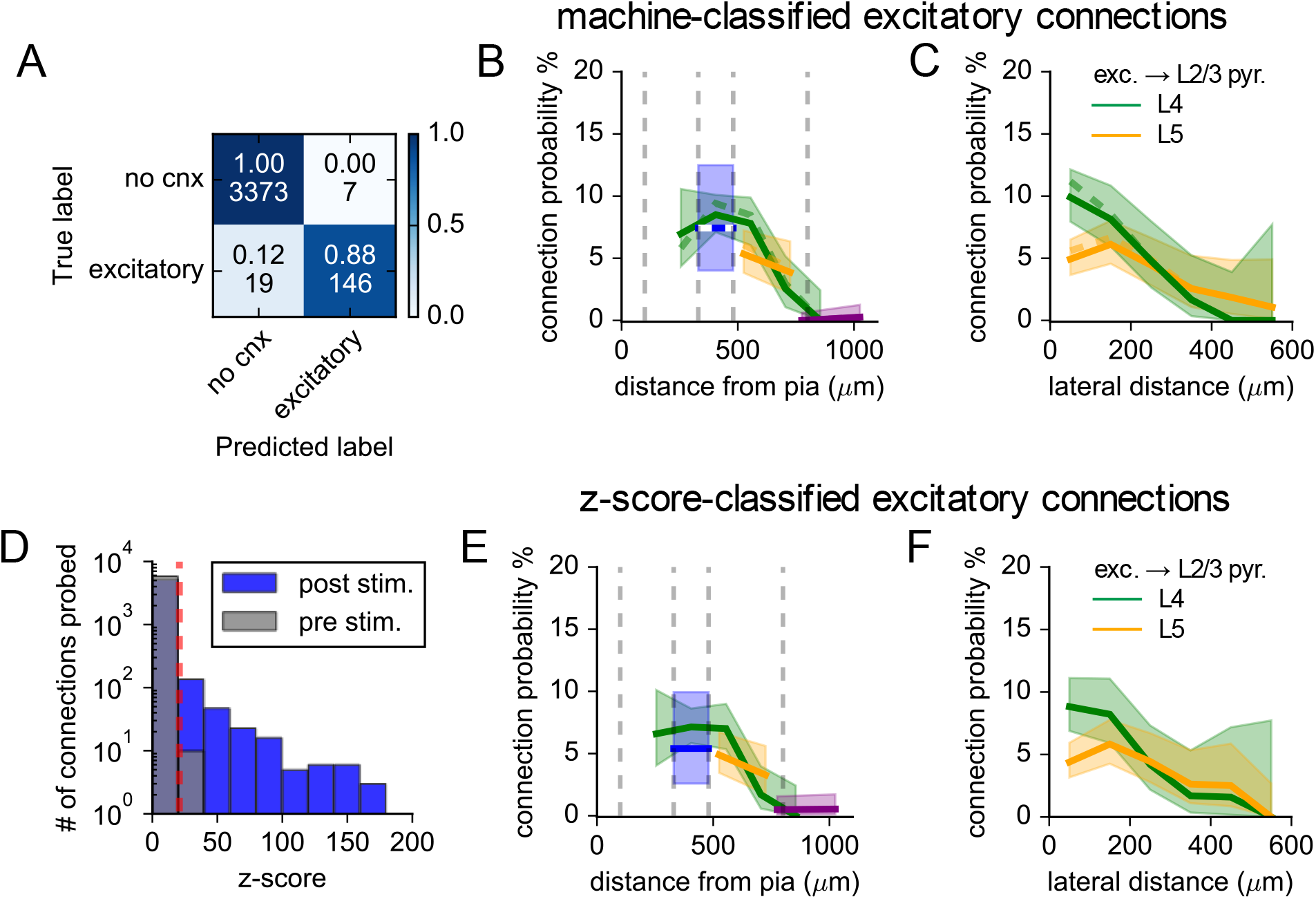
Evaluation of excitatory connectivity. **(A)** Confusion matrix describing the results of a support vector machine tested on 3545 photostimulus responses. The top numbers within each element represent the fraction of human-annotated photoresponses classified by the SVM as containing or not containing an excitatory synaptic response. Bottom values are corresponding counts. **(B)** Connection probability versus the distance of the presynaptic neuron from the pia for Ntsr1- (purple), Rorb- (green), Scnn1a- (blue) and Tlx3- (orange) labeled neurons. **(C)** Connection probability versus the lateral distance between neurons for presynaptic neurons in L4 (green) or L5 (orange). Connection probabilities measured by human annotation are re-plotted from Figure 3 as dashed lines in panels B and C. Solid lines represent connection rates determined by machine classification of the same data. Shading represents 95% confidence intervals. **(D)** Histograms of maximum voltage responses normalized to background variance (z-score) measured 50 ms after photostimulation (post-stimulus) or in a 50 ms window preceding photostimulation (pre - stimulus). Cells were considered connected if the post-stimulus z-score exceeded the 99th percentile of all pre-stimulus z-scores (red dashed line). *E* Connection probability versus distance from the pia (as in panel B) with connected cells classified by z- scores. *F* Connection probability versus the lateral distance between neurons (as in panel C) with connected cells classified by z-scores.

**Figure 4 - Supplement 1.**
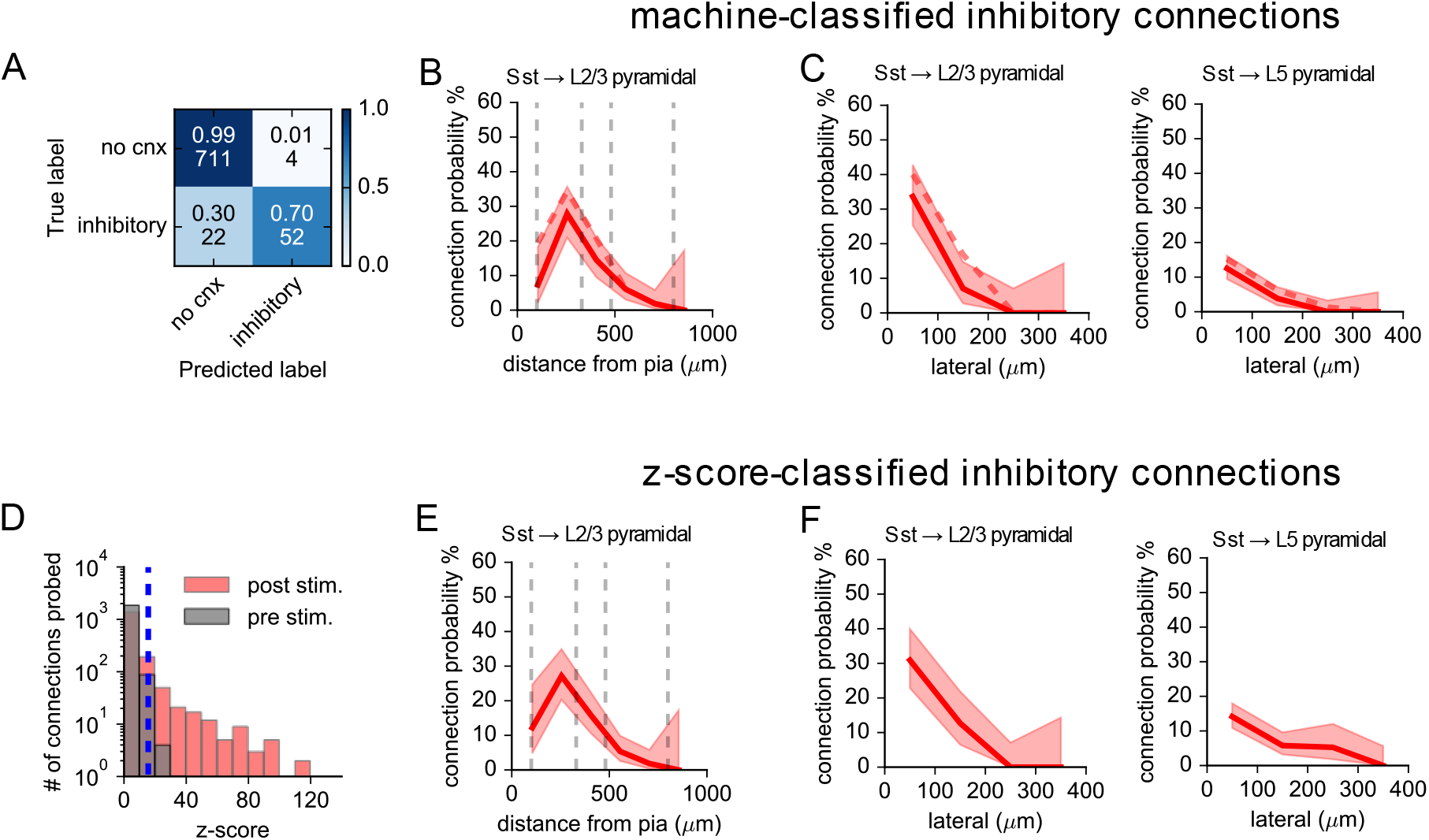
Evaluation of inhibitory connectivity. **(A)** Confusion matrix describing the results of a support vector machine tested on 789 photostimulus responses. The top numbers within each element represent the fraction of human-annotated photoresponses classified by the SVM as containing or not containing an inhibitory synaptic response. Bottom values are corresponding counts. **(B)** Connection probability versus the distance of the presynaptic neuron from the pia for putative Sst to L2/3 pyramidal neuron connections. **(C)** Intralaminar connection probability versus the lateral distance between neurons for Sst and pyramidal cells in L2/3 or in L5. Connection probabilities measured by human annotation are re-plotted from Figure 4 as dashed lines in panels B and C. Solid lines represent connection rates determined by machine classification of the same data. Shading represents 95% confidence intervals. Intralaminar Sst to pyramidal neuron connection probability was significantly lower in L5 than in L2/3 (within 100 μm lateral distance: L2/3: 37/106 probed, 33.6%, L5: 46/368 probed, 12.5%, p=1.3e-6, Fisher’s Exact). **(D)** Histograms of maximum voltage responses normalized to background variance (z-score) measured 50 ms after photostimulation (post-stimulus) or in a 50 ms window preceding photostimulation (pre - stimulus). Cells were considered connected if the post-stimulus z-score exceeded the 99th percentile of all pre-stimulus z-scores (blue dashed line). **(E)** Connection probability versus distance from the pia (as in panel B) with connected cells classified by z-scores. **(F)** Connection probability versus the lateral distance between neurons (as in panel C) with connected cells classified by z-scores. Intralaminar Sst to pyramidal neuron connection probability was significantly lower in L5 than in L2/3 (within 100 μm lateral distance: L2/3: 34/106 probed, 33.6%, L5: 52/368 probed, 12.5%, p=8.4e-5, Fisher’s Exact).

**Figure 4 - Supplement 2.**
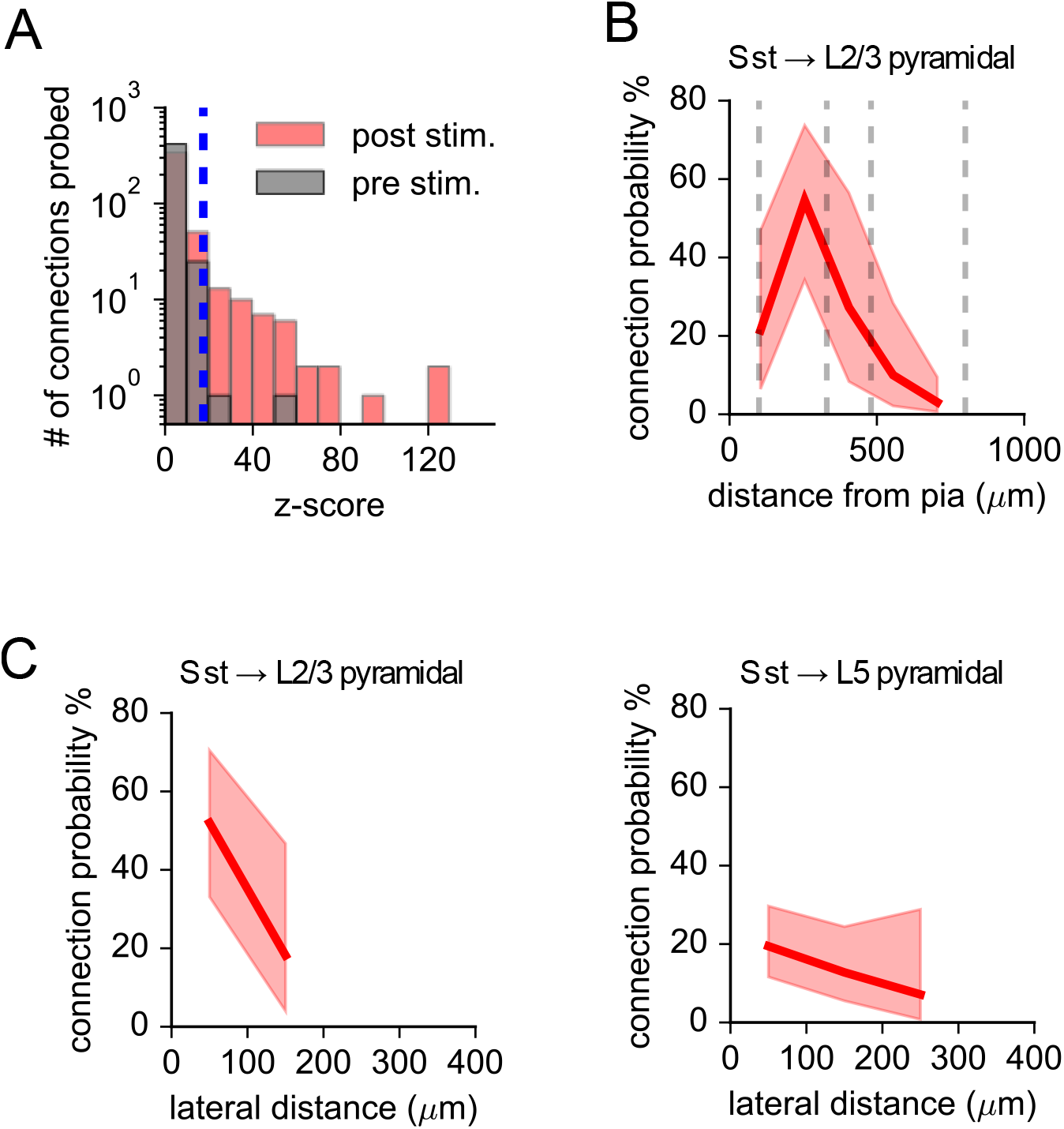
Evaluation of inhibitory connectivity recorded in voltage-clamp. **(A)** Histograms of maximum current responses normalized to background variance (z-score) measured 50 ms after photostimulation (post-stimulus) or in a 50 ms window preceding photostimulation (pre - stimulus). Cells were considered connected if the post-stimulus z-score exceeded the 99th percentile of all pre-stimulus z-scores (blue dashed line). **(B)** Connection probability versus distance from the pia for putative Sst to L2/3 pyramidal neuron connections with connected cells classified by z-scores. **(C)** Intralaminar connection probability versus the lateral distance between neurons for Sst and pyramidal cells in L2/3 or in L5 when connections determined by photostimulus z-score. Shading represents 95% confidence intervals. Intralaminar Sst to pyramidal neuron connection probability was significantly lower in L5 than in L2/3 (within 100 μm lateral distance: L2/3: 13/25 probed, 52%, L5: 13/68 probed, 19.1%, p=0.003, Fisher’s Exact).

**Figure 5 - Supplement 1.**
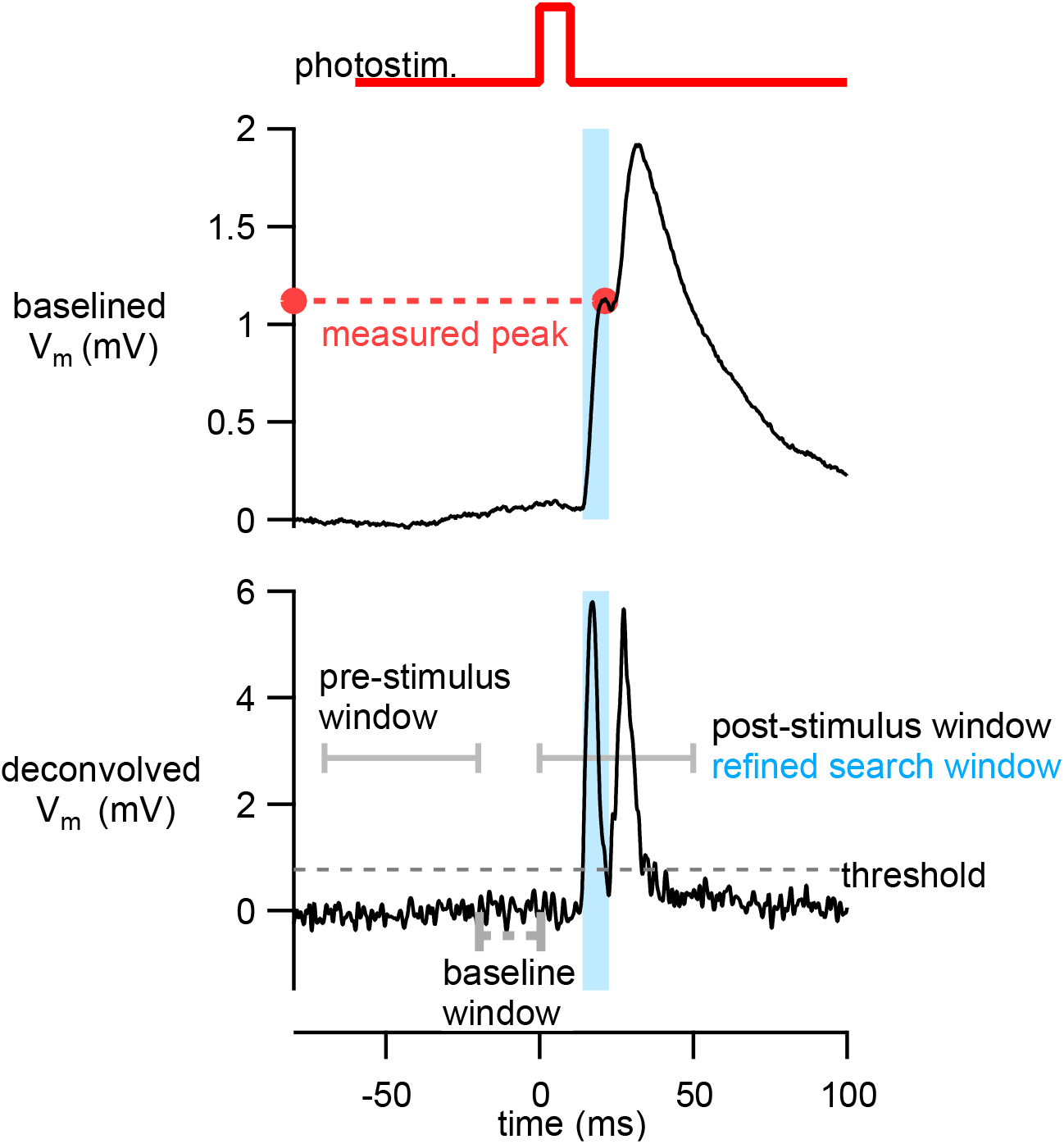
Strategy for measuring first response peak from photoresponses with multiple PSPs. Example of an average membrane potential response from stimulation of a Tlx3 neuron is replotted from Figure 3A. Deconvolved membrane potential was generated using equation 1. An initial response window from 0-50 ms is used to identify timing of PSPs and refine the window in which to measure the peak voltage response in the original trace. See Methods section “Analysis of PSP amplitudes” for further details.

